# Evidence of extensive intraspecific noncoding reshuffling in a 169-kb mitochondrial genome of a basidiomycetous fungus

**DOI:** 10.1101/579870

**Authors:** Hsin-Han Lee, Huei-Mien Ke, Chan-Yi Ivy Lin, Tracy J. Lee, Chia-Lin Chung, Isheng J. Tsai

## Abstract

Comparative genomics of fungal mitochondrial genomes (mitogenomes) have revealed a remarkable pattern of rearrangement between and within major phyla owing to horizontal gene transfer (HGT) and recombination. The role of recombination was exemplified at a finer evolutionary time scale in basidiomycetes group of fungi as they display a diversity of mitochondrial DNA (mtDNA) inheritance patterns. Here, we assembled mitogenomes of six species from the Hymenochaetales order of basidiomycetes and examined 59 mitogenomes from two genetic lineages of *Pyrrhoderma noxium*. Gene order is largely colinear while intergene regions are major determinants of mitogenome size variation. Substantial sequence divergence was found in shared introns consistent with high HGT frequency observed in yeasts, but we also identified a rare case where an intron was retained in five species since speciation. In contrast to the hyperdiversity observed in nuclear genomes of *P. noxium*, mitogenomes’ intraspecific polymorphisms at protein coding sequences are extremely low. Phylogeny based on introns revealed turnover as well as exchange of introns between two lineages. Strikingly, some strains harbor a mosaic origin of introns from both lineages. Analysis of intergenic sequence indicated substantial differences between and within lineages, and an expansion may be ongoing as a result of exchange between distal intergenes. These findings suggest that the evolution in mtDNAs is usually lineage specific but chimeric mitotypes are frequently observed, thus capturing the possible evolutionary processes shaping mitogenomes in a basidiomycete. The large mitogenome sizes reported in various basidiomycetes appear to be a result of interspecific reshuffling of intergenes.

## Introduction

A typical fungal mitochondrial genome (mitogenome) consists of a circular DNA, which encodes 14 mitochondrial-exclusive protein-coding genes, ribosomal RNAs (rRNAs) and transfer RNAs usually in clusters (Férandon, et al. 2013; Aguileta, et al. 2014; Sandor, et al. 2018). Depending on groups, these elements are encoded either on the same strand in ascomycetous or on both in basidiomycetous mitogenomes (Aguileta, et al. 2014). These genomes are highly diversified in size ranging from 19,431 bp in *Schizosaccharomyces pombe* (Bullerwell, et al. 2003) to 235,849 bp in *Rhizoctonia solani* (Losada, et al. 2014). Variations in mitogenome size can be large among species within a genus (Bullerwell, et al. 2003) or among strains of the same species (Wolters, et al. 2015). Presence and expansion of introns in mitogenome protein encoding genes as well as sizable noncoding regions (NCRs) are major determinants of fungal mitogenome size variations (Xu and Wang 2015; Sandor, et al. 2018). Given that the highly variable nature of fungal mitogenome size, characterizing the dynamics of introns and determining the origin and function of these NCRs is critical for understanding the evolution of fungal mitogenome.

Inheritance of mitochondrial DNA (mtDNA) is predominantly maternal in sexual plants and animals (Xu and Wang 2015; Sandor, et al. 2018). In the Fungi kingdom, however, alternative modes of inheritance have been commonly reported during sexual reproduction (Berger and Yaffe 2000; Basse 2010; Wilson and Xu 2012; Xu and Wang 2015). For instance, the model ascomycete yeasts, *Saccharomyces cerevisiae* and *Schizosaccharomyces pombe*, are known to inherit mitochondria biparentally (Wolf, et al. 1976; Basse 2010; Xu and Wang 2015). The zygote contains mtDNA from both parents and subsequently segregated into homoplasmic progenies through the process of syngamy. Like animals and plants, the mating process in filamentous ascomycetes usually initiated with the fusion of two morphologically different gametes, and the mitochondria are typically inherited from the larger gamete (Sandor, et al. 2018). For basidiomycetes, various modes of mitochondrial inheritance may have been involved in generating recombinant mitochondrial genotypes (Xu and He 2015; Xu and Wang 2015; Sandor, et al. 2018).

In fungi, tight regulation of mtDNA transmission is part of molecular mechanisms responsible for sex determination (mating). For model basidiomycetes yeast *Cryptococcus neoformans*, mtDNA is inherited uniparentally from *MAT*a parent butbecomes biparental if the sex-specific homeodomain transcription factor *sxi1α* and *sxi2a* are disrupted (Xu, et al. 2000; Yan and Xu 2003). A closely related species of *C. neoformans*, *C. gattii* exhibited different inheritance patterns depending on strain combinations and environmental factors such as temperature (Voelz, et al. 2013; Zhu, et al. 2013; Wang, et al. 2015). For *U. maydis*, which has a tetrapolar mating system similar to the model mushroom *Coprinopsis cinerea*, the mtDNA inheritance is controlled by *lga2* and *rga2* of uncertain function encoded by *a2* locus (homologous to *C. cinerea matB* locus) (Fedler, et al. 2009).

Strictly selecting mtDNA from only one parent during mating ensures uniparental inheritance. This is the case for model filamentous basidiomycetes such as *C. cinerea* and *Schiozphyllum commune*, where the formation of diploid involves nuclear but not mitochondrial migration, resulting in genetically identical dikaryons with different mitochondria from either parent (Sandor, et al. 2018). Nuclear migration is controlled by *matB* locus encoding peptide pheromones and pheromone receptors (Casselton and Kües 2007), again indicating a strong correlation between mating locus and mitochondria inheritance. Alternatively, the button mushroom *Agaricus bisporus* as well as many basidiomycetes do not undergo nuclear migration and clamp formation; instead, they undergo direct hyphal fusion during mating process. This means different mitochondrial genotypes can be present within hyphal cells, suggesting the opportunity to exchange DNA via mitochondrial recombination. Indeed, *A. bisporus* is known to have uniparental and non-parental mitochondrial inheritance patterns (Hintz, et al. 1988; Jin and Horgen 1993, 1994; de la Bastide and Horgen 2003). Hence, the ephemeral diploid phase may be particularly important in mitochondrial inheritance and diversity (de la Bastide and Horgen 2003; Xu and Wang 2015).

Among the estimated 2.2–3.8 million fungal species (Hawksworth and Lücking 2017), many lineages still lack a formal description or analysis of mitogenomes. One interesting case is the order Hymenochaetales (Agaricomycetes, Basidiomycota), which is dominated by wood decay fungi. Most species in this order are saprotrophic, but some are pathogens involved in major forest incidents since 1971 in different parts of the world (Hepting 1971; Sahashi, et al. 2012). In particular, *Pyrrhoderma noxium* (syn. *Phellinus noxius*) (Zhou, et al. 2018) has a very wide host range, spanning more than 200 broadleaved and coniferous tree species (at least 59 families), and are the causative agents responsible for brown root rot disease in parts of Asia (Akiba, et al. 2015; Chung, et al. 2015). A recent comparative and population genomics study of six genomes from this order and 60 *P. noxium* strains from Taiwan and Japan revealed that *P. noxium* possesses extraordinarily high diversity in its nuclear genomes (Chung, et al. 2017).

*P. noxium* lacks clamp connections (Ann, et al. 2002) and displays a bipolar heterothallic mating system (Chung, et al. 2017) with a highly expanded A locus spanning a ~60-kb region. The availability of whole genome sequences of *P. noxium* populations prompted us to further understand the evolution of mitogenomes in this hypervariable species. In this study, we first assembled and compared the mitogenomes of *P. noxium* and five Hymenochaetales species. We demonstrated that core genes are largely in syntenic across these species while both introns and NCRs constitute the majority of *P. noxium* mtDNA. We then examined mitogenome evolution from two genetically independent lineages of *P. noxium* and discovered frequent rearrangement in NCRs. The main purpose of this study was to elucidate possible mechanisms driving mitogenome enlargement in *P. noxium* at a population level. We provided evidences of frequent mitochondrial exchange possibly in heteroplasmy stage during hyphal fusion and mating resulting in recombinant mitotype and expanding mitogenome.

## Materials and Methods

### *De novo* assembly or reconstruction of four Hymenochaetales mitochondrial genomes

Sequencing data of *Pyrrhoderma noxium*, *Pyrrhoderma lamaoense* (syn. *Phellinus lamaensis*), *Porodaedalea pini* and *Coniferiporia sulphurascens* (syn. *Phellinus sulphurascens*) were described in Chung, et al. (2017). Mitochondrial genomes (mitogenomes) were assembled in 1-6 iterations using Organelle_PBA (v1.0.7; Soorni, et al. 2017) with PacBio reads until sequence circularity was detected. Partial mitochondrial DNA (mtDNA) sequence in the published assembly of each species or conserved protein-coding gene was used as initial reference for the first run. Illumina sequencing data of *Fomitiporia mediterranea* (Floudas, et al. 2012) were obtained from Joint Genome Institute (JGI) database (https://genome.jgi.doe.gov/Fomme1/Fomme1.home.html). The *F. mediterranea* mitogenome was assembled using NOVOplasty (v2.6.7; Dierckxsens, et al. 2017) with *P. pini*’s mitogenome as the seed sequence. Circularity was assessed either by in-built function in the assemblers or subsequently by “check_circularity.pl” script distributed with sprai *de novo* assembly tool (v0.9.9.23; http://zombie.cb.k.u-tokyo.ac.jp/sprai). The consensus of assemblies were improved using pilon (v1.22; Walker, et al. 2014). The complete mitogenome of *Schizopora paradoxa* (Min, et al. 2015) is available in JGI database (https://genome.jgi.doe.gov/Schpa1/Schpa1.home.html) and was retrieved for further annotation and analysis.

### Annotation of mitogenomes

Core genes in mitogenomes were annotated using MFannot web service (http://megasun.bch.umontreal.ca/cgi-bin/mfannot/mfannotInterface.pl) with genetic code table 4. Exon boundaries were further manually curated based on multiple sequence alignment of homologues from closely related species using muscle (v3.8.31; Edgar 2004) and RNA-seq alignments where available. tRNA genes were predicted by MFannot, tRNAscan-SE (v2.0; Lowe and Chan 2016) and aragorn (v1.2.38; Laslett and Canbäck 2004). Only tRNA positions supported by at least two programs were retained for further analysis. Putative open reading frames (ORFs) were predicted by NCBI ORFfinder (v0.4.3; Wheeler, et al. 2003) against exon-masked mitogenomes, and hypothetical proteins were searched against Pfam (Finn, et al. 2016) and NCBI protein nr database (https://www.ncbi.nlm.nih.gov/protein) for evidence of homing endonuclease (HE) and DNA/RNA polymerase (dpo/rpo) genes. Introns were assigned into subgroups using RNAfinder (v1.40; https://github.com/BFL-lab/RNAfinder). Putative homing endonuclease domains in introns were searched by PfamScan (v31; Finn, et al. 2016). Direct and palindromic repeats were searched using Vmatch (v2.3.0; http://www.vmatch.de) with parameters of “-seedlength 30 -l 30 -exdrop 3 -identity 80”. Tandem repeats were identified using Tandem Repeat Finder (v4.09; Benson 1999) with default parameters. Analysis of simple sequence repeats (SSRs) were performed by MISA (v2.0; Thiel, et al. 2003) with default parameters.

### RNA sequencing

The mycelia of *P. noxium* KPN91 were cultured on the PDA agar plates at 25 °C for about 3–5 days. Total RNA was extracted from mycelia homogenized in liquid nitrogen, following with TRIzol™ Reagent (catalog no. 15596018, Thermo Fisher Scientific, Inc.). 200 ng ploy-A enriched RNA was purified from total RNA by using Dynabeads mRNA purification kit (catalog no. 61006, Thermo Fisher Scientific, Inc.), and was used to conduct cDNA library preparation with Direct cDNA Sequencing kit (SQK-DCS108; Oxford Nanopore Technologies). The reads were sequenced with FLO-MIN106 flow cell on a GridION, Oxford Nanopore and then basecalled by dogfish v0.9.3. Illumina RNA-seq reads of four species in this study (*P. noxium*, *P. lamaoense*, *P. pini* and *C. sulphurascens*) were downloaded and aligned to corresponding assemblies using STAR (v2.6.0a; Dobin, et al. 2013). Nanopore cDNA reads (*P. noxium*) were aligned using minimap2 (v2.6; Li 2018). Only reads aligned to mitogenome were retained for further analysis.

### Assembly and annotation of *P. noxium* isolates

The sequencing data of a total of 59 *P. noxium* isolates were described in a previous study (Chung, et al. 2017). All of the mitogenomes were assembled using the same pipeline with *F. mediterranea* as described above except that the seed sequence was *P. noxium* KPN91 mitogenome. Annotation of core genes was searched with query of curated genes of KPN91 using GMAP (v2017-11-15; Wu and Watanabe 2005), followed by multiple sequence alignment using muscle (v3.8.31; Edgar 2004) to identify precise exon-intron boundaries. tRNA annotation was processed as described above. The four (three ‘large’ ranging 6.3–35.4 kb and a small 4.6–8.9 kb) regions were defined and characterised by their frequently rearrangements when compared to KPN91 mtDNA reference (region 1: *nad4*-*rnl*, region 2: *atp8*-*cox1*, region 3: *nad5*-*rns* and region 4s: *cox2*-*cob*).

### Phylogenetic analysis

The core genes phylogeny of six Hymenochatales species was carried out using concatenated amino acid alignments of 15 mitochondrial core genes by muscle (v3.8.31; Edgar 2004), and pruned by trimal strict plus mode (v1.4; Capella-Gutierrez, et al. 2009). A maximum-likelihood phylogeny was computed using iqtree (v1.6.6; Nguyen, et al. 2015) with LG+F+I+G4 model and 100 bootstrap replicates. The same approach was used to produce the intron phylogeny for *P. noxium* isolates, except that ModelFinder (Kalyaanamoorthy, et al. 2017) in iqtree was used to automatically determine the best-fit model. To validate the robustness of mitochondrial core genes phylogeny, four unpublished Hymenochaetales species available on JGI database (*Onnia scaura*, *Phellinus ferrugineofuscus*, *Porodaedalea chrysoloma* and *Porodaedalea niemelaei*) were used to replot the phylogeny.

## Results

### Assembly and annotation of six Hymenochaetales mitochondrial genomes

There are two published mitogenomes in Hymenochaetales species, with genome size ranging from 57.5 kb in *Schizopora paradoxa* (Min, et al. 2015) to 163.4 kb in *Pyrrhoderma noxium* (Chung, et al. 2017). Such large differences in mitogenome sizes prompted us to assemble four more Hymenochaetales species using available Illumina or Pacbio long sequences: *P. lamaoense, P. pini, C. sulphurascens* and *F. mediterranea*. The four Hymenochaetales mitogenomes now range from 45.6 kb to 145.0 kb (table 1), again showing high variations. GC content are low (23.4–34.6%), consistent with the AT rich nature of fungal mitogenomes (Hausner 2003). Remapping of Illumina reads against the mitogenomes showed no heterozygosity, suggesting that mitogenomes are homozygous in these species with no evidence of heteroplasmy. All six species were predicted to contain 14 fungal core protein-coding genes, two ribosomal RNA genes and one ribosomal protein S3 gene (fig. 1, S1 and S2A). 25 tRNAs encoding 20 standard amino acids were shared across 6 species and positioned into 3–5 clusters, with the presence of 1–2 species-specific tRNA. All 17 core genes and tRNA clusters of six species are encoded on the same strand with an exception of a histidine tRNA in *S. paradoxa* (fig. S1E). Interestingly, in four Hymenochaetales species, *nad5* starts (**A**TG) immediately from the last base of adjacent *nad4L* stop codon (TA**A**), which has been observed in various fungal mitochondria (Férandon, et al. 2013; Wang, Zhang, et al. 2018).

**Table 1:**
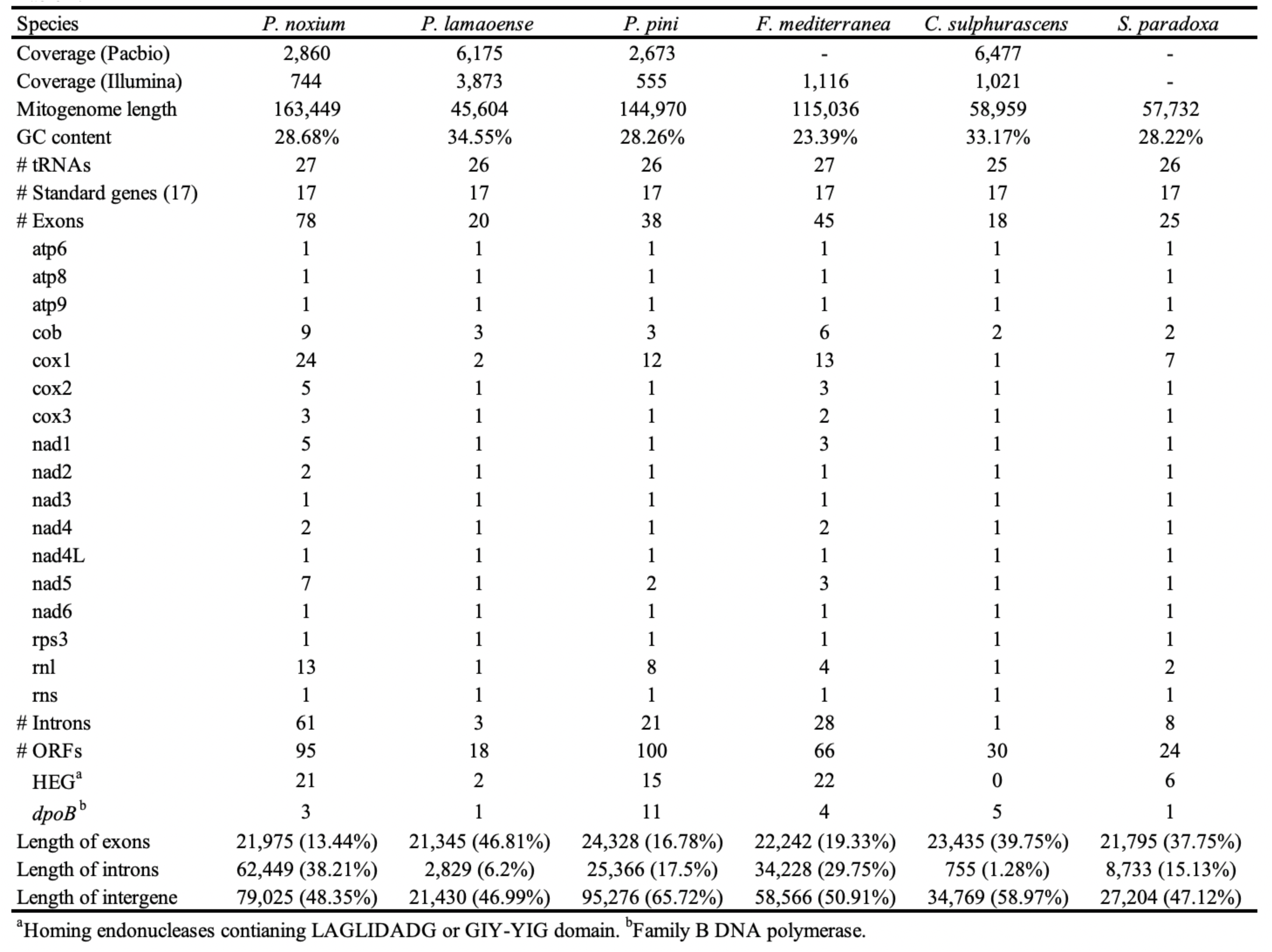
General statistics of six mitogenomes.

**Figure 1:**
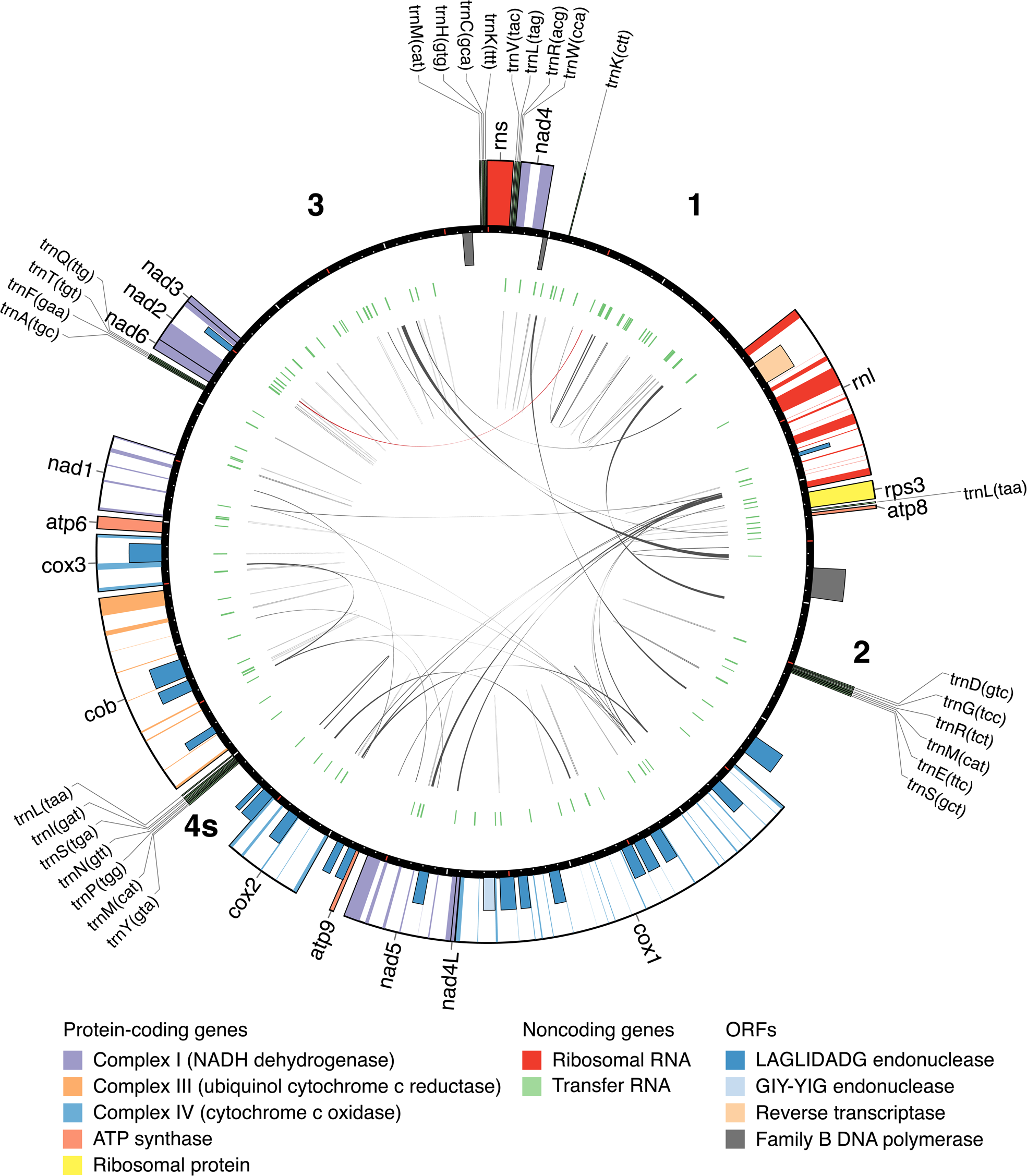
Genetic map of *Pyrrhoderma noxium* KPN91 mitogenome. The first base is defined as the start of *rns* gene. The ticks on the circle are separated by 1 kb. The boxes adhered to the circle are genes and the half-sized boxes are predicted ORFs. The boxes at outside and inside of the circle indicates the genes encoding on positive and negative strand, respectively. Coloured regions are exons. In the inner circle, green tiles represent the tandem repeats and the black and red links represents the direct and palindromic repeats, respectively. The bold numbers indicated the four defined intergeneic regions (1, 2, 3 and 4s). The plot was generated using Circos software (Krzywinski, et al. 2009).

Coding contents are relatively conserved across the six Hymenochaetales species (average 22.5 kb), however, huge variations were observed in their intronic and intergenic sequences. Prediction of open reading frames (ORFs) in intronic and intergenic regions resulted in 18–100 putative protein-coding sequences (table 1 and table S1). Majority of the ORFs did not show homology across various databases. The ORFs with confident protein domain profiles assigned were mostly homing endonuclease (HE, containing LAGLIDADG or GIY-YIG domain) and family B DNA polymerase (*dpoB*). *P. noxium* has the largest mitogenome among the six Hymenochaetales species, with 38.2% introns and 48.4% intergene sequences. The mitogenome of *P. pini* has an intron content of 17.5% and 65.7% intergene region. In contrast, *P. lamaoense* has the most reduced mitogenome with highest percentage of coding content (46.8%). We further investigated whether repetitive sequences contributed to the noncoding content of these mitogenomes. The total repeat content accounted for 2.9%–8.5% of mitogenomes, with only two large palindromic repeats (> 500 bp) observed in *P. pini*. These repeats did not contribute to the majority of noncoding content as only 4.7%–12.2% of intergenic sequences were classified as repetitive. The invertron-like sequences are plasmids integrated into fungal or plant mtDNA, usually consisting of *dpoB* and DNA-dependent RNA polymerase between inverted repeats (Hausner 2012; Mardanov, et al. 2014). Although intact invertron-like sequences were not observed, we found at least one *dpoB* gene in each mitogenome suggesting possible ancient integration events.

### Gene synteny

Species phylogeny inferred from 1,127 orthologous single copy genes placed species of similar genome sizes and primary habits together with *S. paradoxa* being the most diverged clade (Chung, et al. 2017). The mitochondrial phylogeny of six Hymenochaetales was inferred from concatenation of 15 core protein-coding genes (fig. 2). Different from the nuclear phylogeny, *C. sulphurascens* was in the same clade with *P. pini* and *F. mediterranea*. The gene order of 17 core protein, noncoding genes and tRNA clusters were considerably conserved across five of the six Hymenochaetales species. For example, the largest synteny block started from *cox1* and extended to 22 genes (blue box). Greatest difference was found in *S. paradoxa*, which agrees with the nuclear phylogeny (Chung, et al. 2017). In the clade of *P. noxium* and *P. lamaoense*, *nad4* and *rnl* gene were both relocated to downstream of t1 tRNA cluster. Furthermore, the location of t2 tRNA cluster and *rps3*-*atp8* block were swapped in *P. lamaoense*. *P. noxium* and *P. lamaoense* both harboured a tRNA between *nad4* and *rnl*. The two tRNAs shared 88.9% identity in the 72-bp gene region, suggesting accumulation of mutations in a recently gained gene. Boundaries of intergenic spacers were similar between *P. noxium* and *P. lamaoense* but not across other species. Gene duplication events were only observed in t2 in *P. pini* and t5 tRNA clusters in *F. mediterranea*.

**Figure 2:**
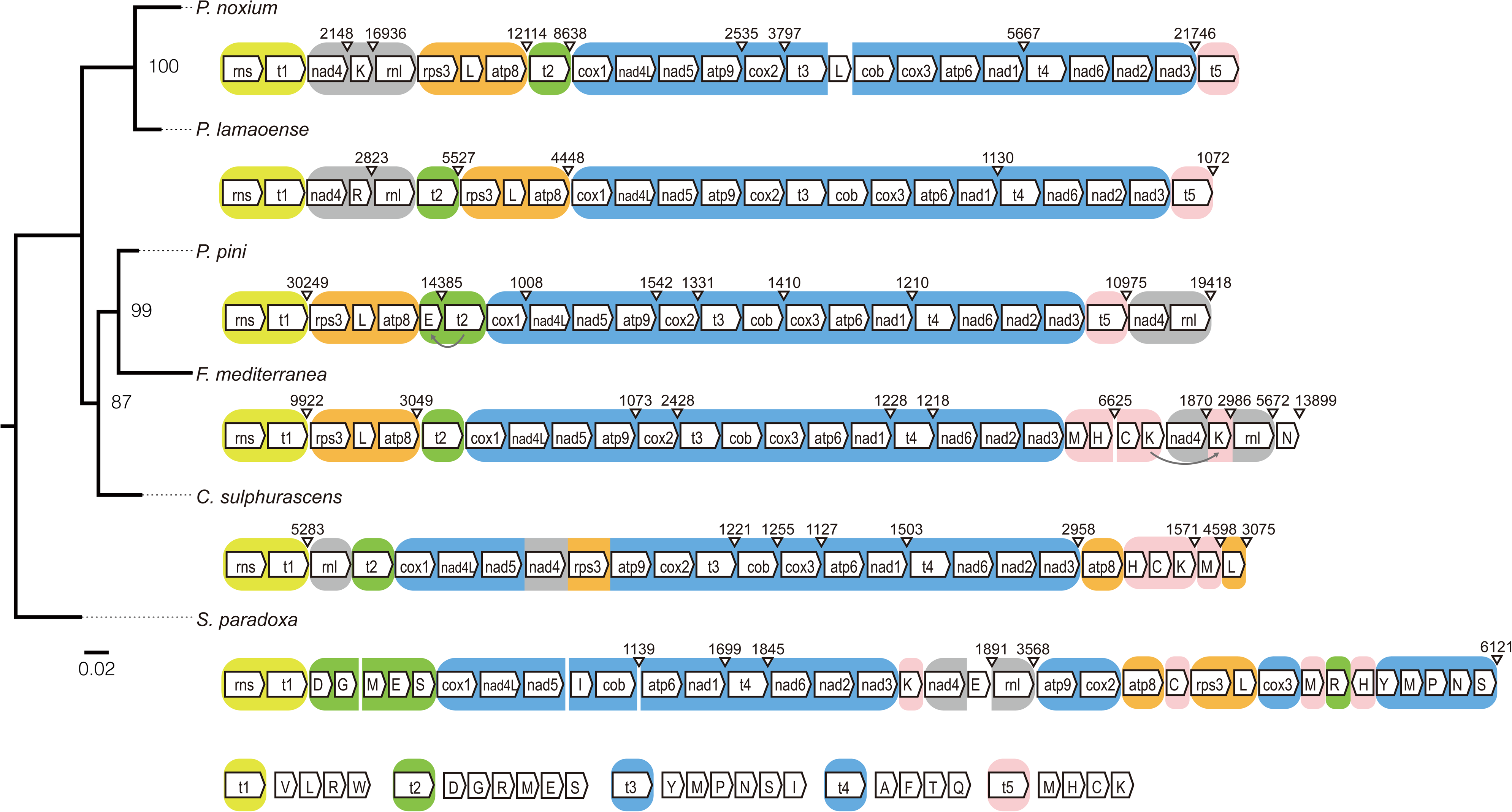
Phylogenetic tree of mitochondrial protein-coding genes and gene synteny in six mitogenomes. The colored boxes represent different conserved gene blocks. The intergene length larger than 1 kb are labeled above the gene wedges. The arrows below the gene wedges indicate gene duplication. The bootstrap values were inferred with 100 replicates.

### Mitochondrial intron dynamics

*P. noxium* mitogenome contains most introns in six species, with 61 introns spreading in 9 genes. In contrast, *C. sulphurascens* mitogenome contains only one intron in *cob* gene. The largest reservoir of introns is *P. noxium cox1* gene, which retains 23 group I introns and a group II intron (17^th^ intron), accounting for 93.5 % of this gene. A total of 122 introns locating in 65 genetic sites were identified in six Hymenochaetales mitogenomes, with 39 sites were shared among species (fig. 3 and table S2). No introns were found in eight genes (*atp6*, *atp8*, *atp9*, *nad3*, *nad4L*, *nad6*, *rns* and *rps3*) whereas the *cob* gene contained at least one intron in every species. The majority of introns were of group I (114/122) and were classified into six subgroups (IA, IA3, IB, IC2, ID and I-derived). Of these, group IB introns accounted for 57.4% of total introns (70/122). Interestingly, the 13^th^ intron of *P. noxium* cox1 gene (cox1_intron13) was predicted to have two separated complete intron cores (IB and ID) suggesting a twintron: an intron inserted by another intron (Deng, et al. 2018).

**Figure 3:**
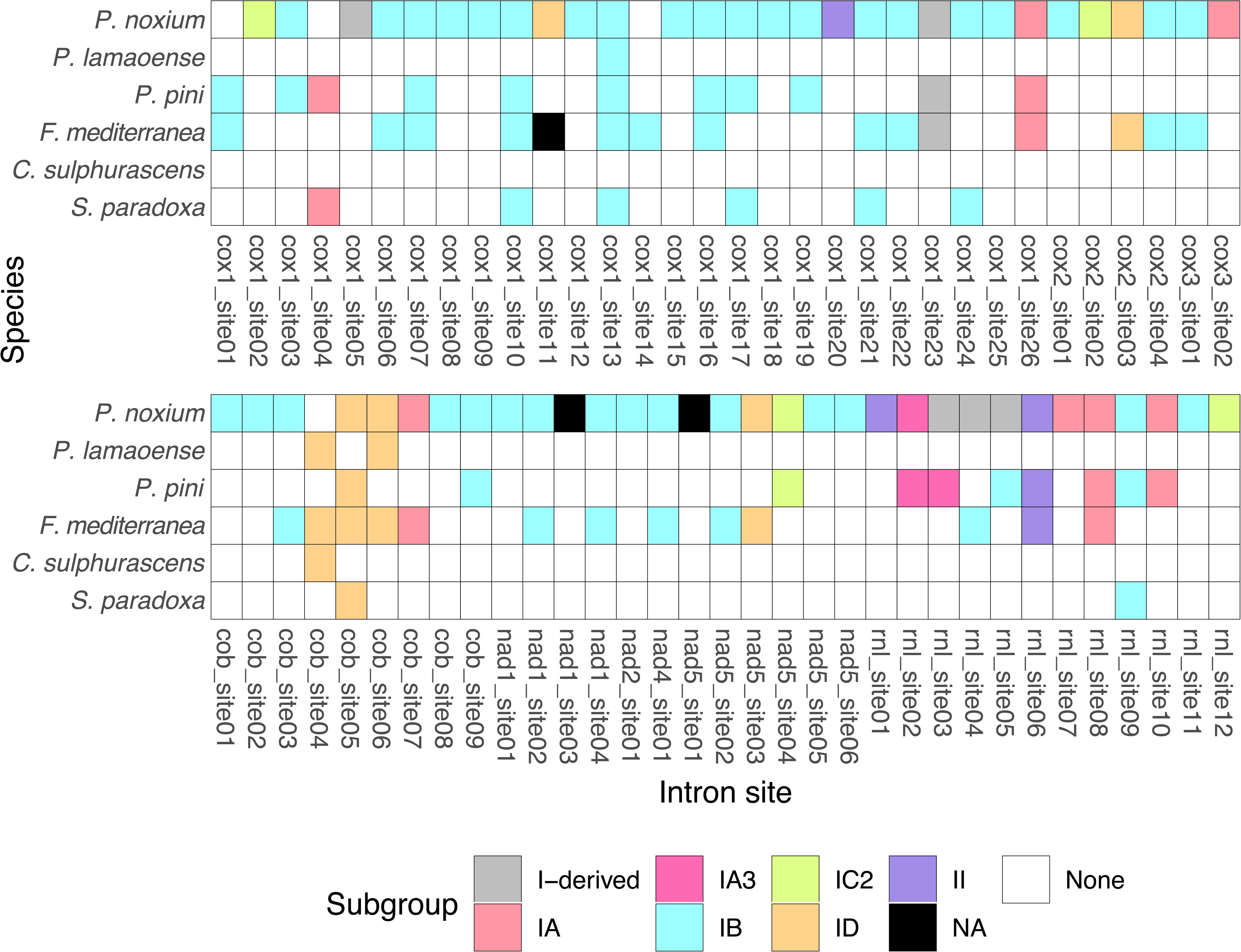
Shared intron positions and intron subgroups in six Hymenochaetales species. Details of intron properties refer to table S2. NA: not assigned to any subgroup.

One possible explanation for the prevalence of introns in these species is the presence of homing endonucleases (HEs), which are highly specific endonucleases playing an important role in intron homing and precise splicing in group I introns (Haugen, et al. 2005; Lang, et al. 2007). We searched for protein domains and ORFs, revealing that 65 introns each containing at least one LAGLIDADG or GIY-YIG domain. Nine of these introns did not have any predicted ORF suggesting loss of HE activity. Conversely, 12 intronic ORFs showed no evidence of HE domain. In addition to group I introns, we found 3 out of 5 group II introns comprising a putative LAGLIDADG domain, which has been reported in several filamentous fungi but showed no activity in enhancing splicing itself. LAGLIDADG domain might be an invaded group I HEG in a group II intron (Toor and Zimmerly 2002; Mullineux, et al. 2010). Only a group II intron (PNOK_rnl_intron01) comprised an intact ORF with complete reverse transcriptase (RT) domain.

Out of 39 intron sites present in multiple species, 34 comprised introns belonging to the same subgroup suggest a preference of insertion (Swithers, et al. 2009). Alternatively, these introns may have already been inserted since their common ancestor; however, this may be less likely as the shared introns showed as low as 74.3% nucleotide identity. This scenario may be more likely in sites where introns are still found in more than three species. There are three such intron sites (cob_site5, cox1_site10 and cox1_site13) and phylogeny indicated two sites (cox1_site10 and cox1_site13) were concordant with the mitochondrial phylogeny, suggesting the intron was present since these species’ common ancestor (fig. S3).

### Population mitogenomics of *P. noxium* – general characteristics

The mitogenomes of additional 58 *P. noxium* isolates from Japan and Taiwan region were assembled into single circular sequences using NOVOPlasty (table S3). The mitogenome size can differ by 22 kb ranging from 146,191 bp of Pn102 to 168,365 bp of Pn120. A strong correlation was observed in intergenic region and mitogenome size (*R*^2^ = 0.987, *P* < 0.001; fig. 4) but not exonic and intronic sequences (*R*^2^ = 0.0213 and *R*^2^ = 0.0203, respectively; fig. 4), suggesting that the expansion of mitochondria was mainly contributed by intergenic regions in *P. noxium*.

**Figure 4:**
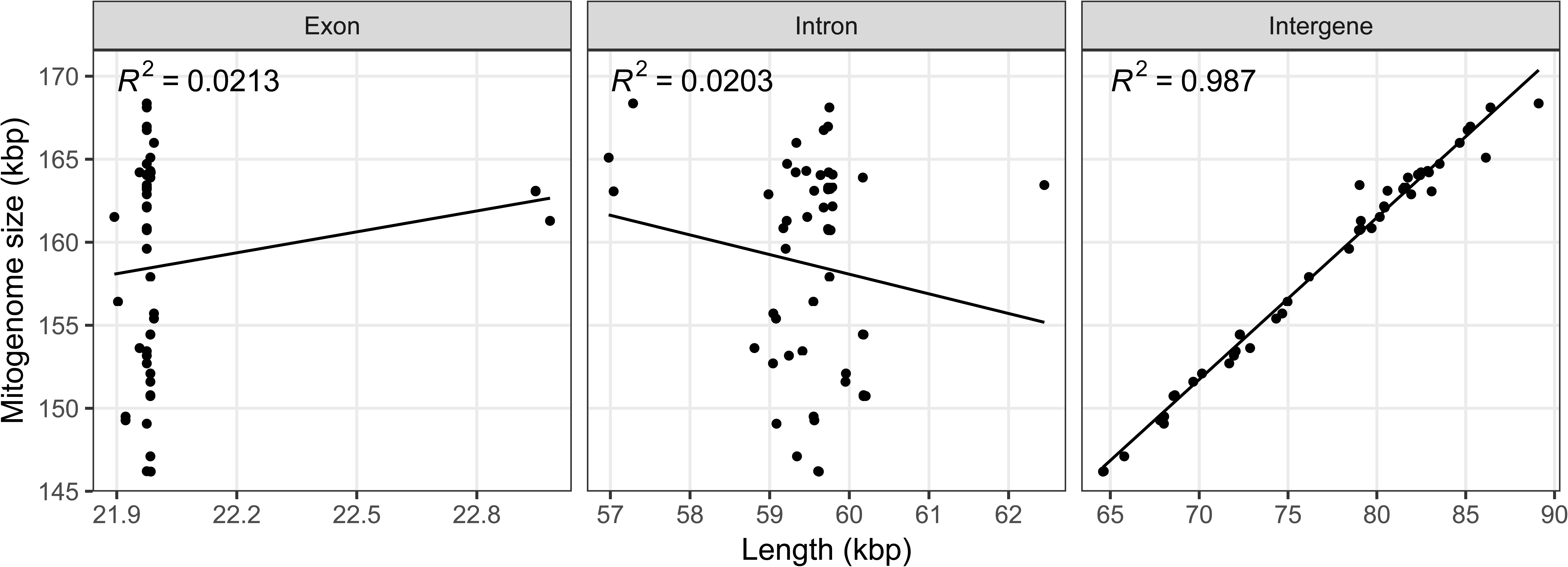
Correlation between mitogenome size and genome properties across 59 *Pyrrhoderma noxium* isolates. Each dot represents an isolate. Details refer to table S3.

### Low intraspecific polymorphism in coding regions

All 17 core genes were present in all isolates. Gene structures were highly conserved except for two protein-coding genes. Stop codon in the majority of isolates in the *cox2* gene was lost in 3 isolates with two possible changes (minor allele 1: KPN57 and Pn315, minor allele 2: Pn040). The prevalent allele encoded for a 765 bp *cox2* gene with 5 exons, and two minor alleles extended 972 bp and 1,008 bp in the last exon, respectively. In *nad2* gene, a tandem repeat [(AGAAACTAA)_3.8–7.8_] located in the second exon varied the length of coding regions from 1,695 bp to 1,731 bp without interrupting the reading frame. Across 4,603 amino acid sites, only 13 and seven were parsimony-informative and singletons, respectively (fig. S4). Number of tRNA ranged from 26– 27. A free-standing tRNA between *nad4* and *rnl* was found to alternatively code for lysine and arginine in these species. Two SNPs were observed in the 72-bp exon region resulting in two different alleles. In 5 isolates (Pn081, Pn102, Pn329, KPN141 and KPN163), a point mutation in the anticodon period turns the lysine tRNA (anticodon: ctt) into an arginine tRNA (anticodon: cct), accompanying the translocation of surrounding sequences (Fig. S7C). The lysine tRNA was lost in 4 isolates (Pn015, Pn035, Pn086 and Pn127) due to large fragment deletion (Fig. S7D), suggesting that it might not be an essential tRNA.

### Frequent gain and loss of introns

Introns range from 57 to 61 in nine intron-containing genes in 59 *P. noxium* isolates (fig. S5). *P. noxium* KPN91, which harboured most introns, had an exclusive group II intron of rnl_intron01 with an intact RT domain, suggesting the intron was recently acquired. Completely loss of introns in some isolates were observed in five intron positions (cox1_intron04, cox1_intron06, cox1_intron07, rnl_intron02 and rnl_intron11). In contrast to the nuclear genome phylogeny, we constructed a phylogeny using the concatenated intronic sequences and revealed three lineages (fig. S6). The majority of Taiwanese and Ryukyu isolates were clustered together, while all eight isolates from Ogasawara island were part of the final group. The result was consistent with the inferences based on nuclear genomes (Chung, et al. 2017). Two Taiwanese isolates were clustered together with the Ogasawara isolates in the intron phylogeny with 100 bootstrap support (fig. S6). We identified three Taiwanese isolates (Pn050, Pn120 and Pn315) as a third lineage, showing a distinct pattern in *cox1* intron 4 to 8 with two introns possessing low identity to other strains and absence of intronic sequences in three sites (fig. 5 and S5).

**Figure 5:**
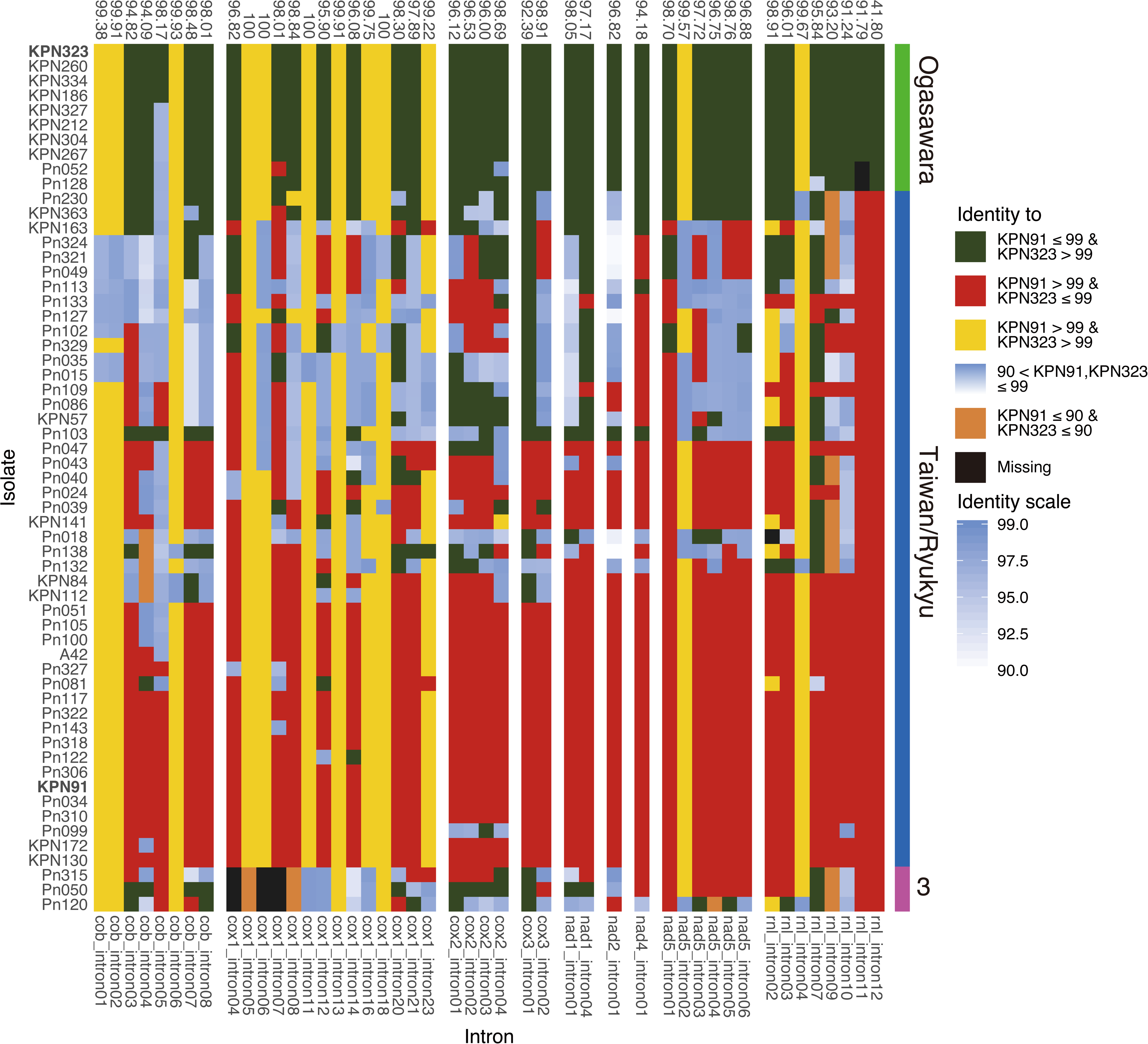
Heatmap of intron identity against KPN91 and KPN323. The intron positions with all identity scores > 99 % against KPN91 are removed. The numbers above the chart indicate the intron identity between KPN91 and KPN323. The tiles on the right of the chart are the lineages defined by intron pattern and intronic phylogeny. The y axis is the order of intronic phylogeny (fig. S6).

To analyse the intron dynamics in these three lineages, we first chose one species from Taiwan/Ryukyu (KPN91) and Ogasarawa (KPN232) lineage as a reference and calculated nucleotide identity of all introns against these references. Intron identity of pairwise comparisons between two references are on average 97.1%, with 11 introns identical to each other. The intron with the lowest pairwise identity was rnl_intron12 with 41.8%, and a closer inspection revealed a unique double sized rnl_intron12. We plotted a heatmap of intron identity according to both references and revealed a mosaic pattern of intron distributions. With the double sized rnl_intron12, the introns of the two Taiwanese isolates that were clustered together to the Ogasawara lineage in the intron phylogeny resemble the introns of KPN323 more than that of KPN91 (Fig. 5). In contrast, strains belonging to the Taiwan/Ryukyu lineage have revealed striking differences amongst strains (fig. 5). Three modes were found: i) strains with introns that completely resemble the Taiwanese/Ryukyu reference, ii) strains with mosaic intron patterns that resemble either Taiwan/Ryukyu or Ogasawara references, iii) strains with introns that were equally dissimilar to both references. These observations suggest that extensive rearrangement of introns have occurred in the Taiwan/Ryukyu lineage but less so in the Ogasawara lineage.

### Evidence of extensive shuffling between large intergene regions separated by gene clusters

To understand the causes of large intergene that are present in *P. noxium*, we first defined three large (region 1: *nad4*–*rnl* 11.3–25.9 kb, region 2: *atp8*–*cox1* 15.3–35.4 kb and region 3: *nad3*–*rns* 6.3–24.9 kb) and a small (region 4s: *cox2*-*cob* 4.6– 8.9 kb) intergenic regions. A close species *P. lamaoense* had only 11.3 kb in the regions we defined, comparing to an average of 66.2 kb in *P. noxium*. MtDNA alignment also showed no similarity with other species in this study, suggesting that these intergenic regions are *P. noxium*-specific (fig. S2B).

Frequent intra- and inter-rearrangements between four regions were observed. We first sought if a correlation exists between the intron phylogeny and intergene. However, in the two distinct clades defined by the intron phylogeny, similar synteny was observed in isolates belonging to different clades (fig. S7A, B). To investigate the effect of rearrangement in mitogenome evolution, we took KPN91 as a reference and sought if any rearrangement event occurred in the four defined intergenic regions (fig. S8). The 10 isolates in Ogasawara lineage had similar rearrangement pattern, and the intergene sizes were significantly shorter than that of Taiwan/Ryukyu lineage isolates (*P* < 0.01; fig. 6). Isolates from the Taiwan/Ryukyu were subsequently classified into two groups based on intergene rearrangement pattern against KPN91 (fig. S8). They were also significantly different in length (*P* < 0.001; fig. 6). The length of intergenic spacers in the isolates with rearrangement tended to be smaller than that of the isolates without rearrangement, resulting in a median difference of ~9.7 kb. Additionally, 5 isolates with intra-rearrangement in region 1 were accompanied with the translocation of a free-standing tRNA between *nad4* and *rnl*, suggesting that it was potentially a foreign gene (fig. S7C and D).

**Figure 6:**
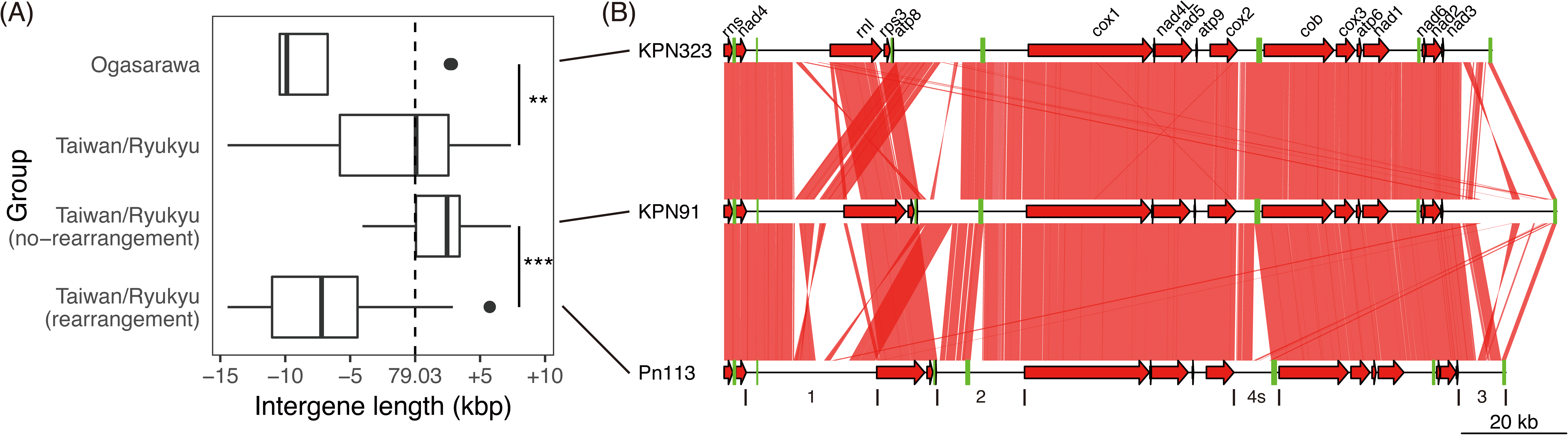
Intergene length distribution and rearrangement events between Ogasawara and Taiwan/Ryukyu lineages. (A) The intergene length between lineages and rearranged or non-rearranged intergene against KPN91 (fig. S8) are significantly different (**: *P* < 0.01, ***: *P* < 0.001). The dotted line represents KPN91 intergene length. (B) Rearrangement events in four defined regions between lineages in (A). Red arrows are mitochondrial core genes and green tiles are tRNA genes. Red links between mitogenomes are aligned regions with identity > 80%. The intergene boundaries are labelled under Pn113. The plot was generated with genoplotR package in R.

## Discussion

In our previous study, we reported four Hymenochaetales genomes and revealed the nature of hyperdiversity of nuclear genomes in two lineages of *P. noxium* in Taiwan and Japan areas (Chung, et al. 2015; Chung, et al. 2017). To understand the mitochondrial genomics in Hymenochaetales, we analysed six mitogenomes in this order. All mitogenomes can be circularized, indicating the high quality of these assemblies. All fungal mitochondrial core genes are present in these species, and the synteny are largely conserved according to their phylogenetic relationships. The overlapping start and stop codons of *nad* genes have been observed in not only fungi but also animals (Franco, et al. 2017; Wang, Shi, et al. 2018), presumably because of the mRNA transcription of these genes have remained coupled across independent lineages. Only tRNAs were found duplicated and lost, presumably due to the preference of mitochondrial recombination in these regions (Fritsch, et al. 2014).

The six mitogenomes varied by at most 3.6-fold in length, from 45.6 kb in *P. lamaoense* to 163.4 kb in *P. noxium*. In Organelle Genome Resources NCBI, only six species are larger than *P. noxium* (Losada, et al. 2014; Mardanov, et al. 2014; Kanzi, et al. 2016; Nowrousian 2016). Gain of introns have been depicted as the main constituents of the enlargement of fungal mitogenomes (Losada, et al. 2014; Mardanov, et al. 2014; Salavirta, et al. 2014; Kanzi, et al. 2016). In the case of Hymenochaetales, three out of the six species hold an impressively high number of introns (table 1). In particular, *cox1* harbours up to 23 introns in these species which is consistent to previous surveys in fungal mitogenomes as the most intron-containing gene.

Horizontal gene transfer (HGT) event have been implicated as the main causes and origins of intron gains in fungi (Hausner 2012; Wu, et al. 2015). We determined 39 out of 65 shared intron positions, however, only 14 and three intron sites were shared in more than two or three species, respectively. From these sites, we found only two examples of shared intron exhibiting same phylogenetic relationship as the species tree, implying an ancient transposition event. Additional evidence from comparative analysis of multiple species within an order will help reveal whether the retention of such introns has any selective advantage.

Intergenic regions constitute the majority of mitogenomes in all species in this study, consistent with previous findings in yeast mitogenomes where intergenic sequences played a role in genome expansion (Xiao, et al. 2017). The initial expansion of intergenic regions may be contributed by the integration of plasmid-like sequences since the few numbers of repetitive elements (Hausner 2003; Férandon, et al. 2013). Unlike the introns which need to be spliced precisely for mature mRNA, the integrated plasmid sequences are easily degenerated overtime due to the absence of selection pressure (Hausner 2003; Swithers, et al. 2009). Integration of invertron plasmid in mtDNA has been reported in many fungi species including *Moniliophthora perniciosa*, *Agaricus bisporus*, and *Sclerotinia borealis* (Formighieri, et al. 2008; Férandon, et al. 2013; Mardanov, et al. 2014). We found at least one family B DNA polymerase located in intergenic regions in all six species (table 1), but no putative RNA polymerase or invertron-like repetitive elements, suggesting degenerated integrated-plasmid.

Our previous investigation on population-scale sequencing of *P. noxium* nuclear genomes suggests that it is a hypervariable species. Hence, one of the aims of this study was to investigate the evolution history of *P. noxium* from the mitochondrial perspective. Surprisingly, the analysis of 59 isolates revealed different pace of evolution across coding, introns and intergenic regions. The coding regions unusually harbor low variation across the two lineages, suggesting strong purifying selection, i.e., arising substitutions were quickly purged from the population. Considering the much higher mutation rate in mitogenome than the nuclear genome (Jung, et al. 2012), additional regulatory mechanisms may exist in hypervariable species to reduce potential detrimental effects from the replication of mitogenome.

A previous investigation of intraspecific variations on three species of yeast determined introns contributed the highest variance in genome size (Xiao, et al. 2017). Our intron analyses of two lineages of *P. noxium* revealed high similarity but extensive rearrangements of introns among strains of different clades. Based on current knowledge in ascomycetes, there are two possible scenarios that can account for such observations in a basidiomycete during hyphal fusion between strains of two mitotypes: intron homing or homologous recombination (Haugen, et al. 2005; Wu, et al. 2015; Leducq, et al. 2017). We observed few gain and loss of introns at population level so recombination may be a more predominant process, which has been documented in various fungal species (Leducq, et al. 2017). Unfortunately, high similarity of intron sequences impedes further recombination-detection based analyses. Future investigations should focus on determination of mitochondrial inheritance mode and the frequency and mechanisms involved in intron rearrangements using artificial hybrids.

For the first time in our investigation, large variations at intergenic regions are documented at an intraspecies level, even though it has been commonly observed between closely related fungal species (Burger, et al. 2003; Aguileta, et al. 2014; Himmelstrand, et al. 2014). We have observed contrasting differences in intergene length between strains of different lineages as well as within a lineage. These findings suggest that, when compared to closely related species, intergene variations are currently driving intraspecific mitogenomic variation in *P. noxium* and may eventually lead to larger intergene variations. Intriguingly, repeats do not seem to play a major role in shaping such variations and it remains to be determined whether homologous recombination will lead to extension of intergenic sequences.

In conclusion, we have revealed for the first time that intergenic regions play a major role in shaping and extending the mtDNA in Basdiomycetes at inter- and intraspecies level. The extreme contrast between nuclear genome and coding region of mitogenome suggests strong purifying selection acting on these genes. Basidiomycetes can be excellent models in understanding mtDNA inheritance. A comprehensive sampling and detailed comparison across mitogenome will help delineate the relationship between mating and mitochondrial diversity.

## Authors contribution

I.J.T conceived the study. H.M.K, C.Y.I.L, T.J.L and C.L.C. established the strains collection, carried out experiments and sequencing. H.H.L carried out comparative analysis. I.J.T. and H.H.L wrote the manuscript.

## Data availability

The complete mitochondrial genomes generated in this study have been deposited in NCBI and will be publicized under accession MK623257-MK623261 once internal processing are completed.

---

Table S1: Putative intronic and intergenic ORFs and pfam annotations.

Table S2: Details of intron properties and HMM domain prediction. The protein domains were predicted against intron sequences directly, hence may not locate in ORF.

Table S3: Length of different mitogenome properties of 59 *Pyrrhoderma noxious* isolates.

## References

Aguileta G, de Vienne DM, Ross ON, Hood ME, Giraud T, Petit E, Gabaldón T. 2014. High variability of mitochondrial gene order among fungi. Genome Biology and Evolution 6:451–465.

Akiba M, Ota Y, Tsai IJ, Hattori T, Sahashi N, Kikuchi T. 2015. Genetic differentiation and spatial structure of *Phellinus noxius*, the causal agent of brown root rot of woody plants in Japan. Plos One 10:e0141792.

Ann P-J, Chang T-T, Ko W-H. 2002. *Phellinus noxius* brown root rot of fruit and ornamental trees in Taiwan. Plant Disease 86:820–826.

Basse CW. 2010. Mitochondrial inheritance in fungi. Current opinion in microbiology 13:712–719.

Benson G. 1999. Tandem repeats finder: a program to analyze DNA sequences. Nucleic Acids Research 27:573–580.

Berger KH, Yaffe MP. 2000. Mitochondrial DNA inheritance in Saccharomyces cerevisiae. Trends in microbiology 8:508–513.

Bullerwell CE, Leigh J, Forget L, Lang BF. 2003. A comparison of three fission yeast mitochondrial genomes. Nucleic Acids Research 31:759–768.

Burger G, Gray MW, Franz Lang B. 2003. Mitochondrial genomes: anything goes. Trends in genetics: TIG 19:709–716.

Capella-Gutierrez S, Silla-Martinez JM, Gabaldon T. 2009. trimAl: a tool for automated alignment trimming in large-scale phylogenetic analyses. Bioinformatics 25:1972–1973.

Casselton LA, Kües U. 2007. The origin of multiple mating types in the model mushrooms *Coprinopsis cinerea* and *Schizophyllum commune*. In: Heitman J, Kronstad JW, Taylor JW, Casselton LA, editors: American Society of Microbiology. p. 283–300.

Chung C-L, Huang S-Y, Huang Y-C, Tzean S-S, Ann P-J, Tsai J-N, Yang C-C, Lee H-H, Huang T-W, Huang H-Y, et al. 2015. The genetic structure of *Phellinus noxius* and dissemination pattern of brown root rot disease in Taiwan. Plos One 10:e0139445.

Chung C-L, Lee TJ, Akiba M, Lee H-H, Kuo T-H, Liu D, Ke H-M, Yokoi T, Roa MB, Lu M-YJ, et al. 2017. Comparative and population genomic landscape of *Phellinus noxius*: A hypervariable fungus causing root rot in trees. Molecular Ecology 10:e0141792–0141716.

de la Bastide PY, Horgen PA. 2003. Mitochondrial inheritance and the detection of non-parental mitochondrial DNA haplotypes in crosses of *Agaricus* bisporus homokaryons. Fungal Genetics and Biology 38:333–342.

Deng Y, Hsiang T, Li S, Lin L, Wang Q, Chen Q, Xie B, Ming R. 2018. Comparison of the mitochondrial genome sequences of six *Annulohypoxylon stygium* isolates suggests short fragment insertions as a potential factor leading to larger genomic size. Frontiers in microbiology 9:380.

Dierckxsens N, Mardulyn P, Smits G. 2017. NOVOPlasty: de novo assembly of organelle genomes from whole genome data. Nucleic Acids Research 45:e18.

Dobin A, Davis CA, Schlesinger F, Drenkow J, Zaleski C, Jha S, Batut P, Chaisson M, Gingeras TR. 2013. STAR: ultrafast universal RNA-seq aligner. Bioinformatics 29:15–21.

Edgar RC. 2004. MUSCLE: a multiple sequence alignment method with reduced time and space complexity. BMC bioinformatics 5:113.

Fedler M, Luh K-S, Stelter K, Nieto-Jacobo F, Basse CW. 2009. The *a2* mating-type locus genes *lga2* and *rga2* direct uniparental mitochondrial DNA (mtDNA) inheritance and constrain mtDNA recombination during sexual development of *Ustilago maydis*. Genetics 181:847–860.

Férandon C, Xu J, Barroso G. 2013. The 135 kbp mitochondrial genome of *Agaricus bisporus* is the largest known eukaryotic reservoir of group I introns and plasmid-related sequences. Fungal Genetics and Biology 55:85–91.

Finn RD, Coggill P, Eberhardt RY, Eddy SR, Mistry J, Mitchell AL, Potter SC, Punta M, Qureshi M, Sangrador-Vegas A, et al. 2016. The Pfam protein families database: towards a more sustainable future. Nucleic Acids Research 44:D279–D285.

Floudas D, Binder M, Riley R, Barry K, Blanchette RA, Henrissat B, Martinez AT, Otillar R, Spatafora JW, Yadav JS, et al. 2012. The Paleozoic origin of enzymatic lignin decomposition reconstructed from 31 fungal genomes. Science 336:1715–1719.

Formighieri EF, Tiburcio RA, Armas ED, Medrano FJ, Shimo H, Carels N, Góes-Neto A, Cotomacci C, Carazzolle MF, Sardinha-Pinto N, et al. 2008. The mitochondrial genome of the phytopathogenic basidiomycete *Moniliophthora perniciosa* is 109 kb in size and contains a stable integrated plasmid. Mycological Research 112:1136–1152.

Franco MEE, López SMY, Medina R, Lucentini CG, Troncozo MI, Pastorino GN, Saparrat MCN, Balatti PA. 2017. The mitochondrial genome of the plant-pathogenic fungus *Stemphylium lycopersici* uncovers a dynamic structure due to repetitive and mobile elements. Plos One 12:e0185545.

Fritsch ES, Chabbert CD, Klaus B, Steinmetz LM. 2014. A genome-wide map of mitochondrial DNA recombination in yeast. Genetics 198:755–771.

Haugen P, Simon DM, Bhattacharya D. 2005. The natural history of group I introns. Trends in genetics: TIG 21:111–119.

Hausner G. 2003. Fungal Mitochondrial Genomes, Plasmids and Introns. In: Arora DK, Khachatourians GG, editors: Elsevier. p. 101–131.

Hausner G. 2012. Introns, Mobile Elements, and Plasmids. In. Berlin, Heidelberg: Springer Berlin Heidelberg. p. 329–357.

Hawksworth DL, Lücking R. 2017. Fungal diversity revisited: 2.2 to 3.8 million species. Microbiology spectrum 5.

Hepting GH. 1971. Diseases of forest and shade trees of the United States. Washington, DC: U.S. Department of Agriculture Forest Service.

Himmelstrand K, Olson Å, Durling MB, Karlsson M, Stenlid J. 2014. Intronic and plasmid-derived regions contribute to the large mitochondrial genome sizes of Agaricomycetes. Current Genetics 60:303–313.

Hintz W, Anderson JB, Horgen PA. 1988. Nuclear migration and mitochondrial inheritance in the mushroom *Agaricus bitorquis*. Genetics 119:35–41.

Jin T, Horgen PA. 1993. Further characterization of a large inverted repeat in the mitochondrial genomes of *Agaricus bisporus* (= *A. brunnescens*) and related species. Current Genetics 23:228–233.

Jin T, Horgen PA. 1994. Uniparental mitochondrial transmission in the cultivated button mushroom, *Agaricus bisporus*. Applied and environmental microbiology 60:4456–4460.

Jung PP, Friedrich A, Reisser C, Hou J, Schacherer J. 2012. Mitochondrial genome evolution in a single protoploid yeast species. G3-Genes Genomes Genetics 2:1103–1111.

Kalyaanamoorthy S, Minh BQ, Wong TKF, von Haeseler A, Jermiin LS. 2017. ModelFinder: fast model selection for accurate phylogenetic estimates. Nature Methods 14:587–589.

Kanzi AM, Wingfield BD, Steenkamp ET, Naidoo S, van der Merwe NA. 2016. Intron derived size polymorphism in the mitochondrial genomes of closely related *Chrysoporthe species*. Plos One 11:e0156104.

Krzywinski MI, Schein JE, Birol I, Connors J, Gascoyne R, Horsman D, Jones SJ, Marra MA. 2009. Circos: An information aesthetic for comparative genomics. Genome research 19:1639–1645.

Lang BF, Laforest M-J, Burger G. 2007. Mitochondrial introns: a critical view. Trends in genetics: TIG 23:119–125.

Laslett D, Canbäck B. 2004. ARAGORN, a program to detect tRNA genes and tmRNA genes in nucleotide sequences. Nucleic Acids Research 32:11–16.

Leducq J-B, Henault M, Charron G, Nielly-Thibault L, Terrat Y, Fiumera HL, Shapiro BJ, Landry CR. 2017. Mitochondrial recombination and introgression during speciation by hybridization. Molecular Biology and Evolution 34:1947–1959.

Li H. 2018. Minimap2: pairwise alignment for nucleotide sequences. Bioinformatics 34:3094–3100.

Losada L, Pakala SB, Fedorova ND, Joardar V, Shabalina SA, Hostetler J, Pakala SM, Zafar N, Thomas E, Carres MR, et al. 2014. Mobile elements and mitochondrial genome expansion in the soil fungus and potato pathogen Rhizoctonia solani AG-3. FEMS Microbiology Letters 352:165–173.

Lowe TM, Chan PP. 2016. tRNAscan-SE On-line: integrating search and context for analysis of transfer RNA genes. Nucleic Acids Research 44:W54–57.

Mardanov AV, Beletsky AV, Kadnikov VV, Ignatov AN, Ravin NV. 2014. The 203 kbp mitochondrial genome of the phytopathogenic fungus *Sclerotinia borealis* reveals multiple invasions of introns and genomic duplications. Plos One 9:e107536–107511.

Min B, Park H, Jang Y, Kim J-J, Kim KH, Pangilinan J, Lipzen A, Riley R, Grigoriev IV, Spatafora JW, et al. 2015. Genome sequence of a white rot fungus *Schizopora paradoxa* KUC8140 for wood decay and mycoremediation. Journal of biotechnology 211:42–43.

Mullineux S-T, Costa M, Bassi GS, Michel F, Hausner G. 2010. A group II intron encodes a functional LAGLIDADG homing endonuclease and self-splices under moderate temperature and ionic conditions. Rna 16:1818–1831.

Nguyen L-T, Schmidt HA, von Haeseler A, Minh BQ. 2015. IQ-TREE: a fast and effective stochastic algorithm for estimating maximum-likelihood phylogenies. Molecular Biology and Evolution 32:268–274.

Nowrousian M. 2016. Complete mitochondrial genome sequence of the Pezizomycete *Pyronema confluens*. Genome announcements 4:e107536.

Sahashi N, Akiba M, Ishihara M, Ota Y, Kanzaki N. 2012. Brown root rot of trees caused by *Phellinus noxius* in the Ryukyu Islands, subtropical areas of Japan. Forest Pathology 42:353–361.

Salavirta H, Oksanen I, Kuuskeri J, Mäkelä M, Laine P, Paulin L, Lundell T. 2014. Mitochondrial genome of *Phlebia radiata* is the second largest (156 kbp) among fungi and features signs of genome flexibility and recent recombination events. Plos One 9:e97141.

Sandor S, Zhang Y, Xu J. 2018. Fungal mitochondrial genomes and genetic polymorphisms. Applied Microbiology and Biotechnology 102:9433–9448.

Soorni A, Haak D, Zaitlin D, Bombarely A. 2017. Organelle_PBA, a pipeline for assembling chloroplast and mitochondrial genomes from PacBio DNA sequencing data. BMC Genomics 18:49.

Swithers KS, Senejani AG, Fournier GP, Gogarten JP. 2009. Conservation of intron and intein insertion sites: implications for life histories of parasitic genetic elements. BMC Evolutionary Biology 9:303.

Thiel T, Michalek W, Varshney R, Graner A. 2003. Exploiting EST databases for the development and characterization of gene-derived SSR-markers in barley (*Hordeum vulgare*). Theoretical and Applied Genetics 106:411–422.

Toor N, Zimmerly S. 2002. Identification of a family of group II introns encoding LAGLIDADG ORFs typical of group I introns. Rna 8:1373–1377.

Voelz K, Ma H, Phadke S, Byrnes EJ, Zhu P, Mueller O, Farrer RA, Henk DA, Lewit Y, Hsueh Y-P, et al. 2013. Transmission of hypervirulence traits via sexual reproduction within and between lineages of the human fungal pathogen *Cryptococcus gattii*. PLoS Genetics 9:e1003771.

Walker BJ, Abeel T, Shea T, Priest M, Abouelliel A, Sakthikumar S, Cuomo CA, Zeng Q, Wortman J, Young SK, et al. 2014. Pilon: an integrated tool for comprehensive microbial variant detection and genome assembly improvement. Plos One 9:e112963.

Wang L, Zhang S, Li J-H, Zhang Y-J. 2018. Mitochondrial genome, comparative analysis and evolutionary insights into the entomopathogenic fungus Hirsutella thompsonii. Environmental Microbiology 7:916–939.

Wang Z, Shi X, Tao Y, Wu Q, Bai Y, Guo H, Tang D. 2018. The complete mitochondrial genome of Parasesarma pictum (Brachyura: Grapsoidea: Sesarmidae) and comparison with other Brachyuran crabs. Genomics.

Wang Z, Wilson A, Xu J. 2015. Mitochondrial DNA inheritance in the human fungal pathogen *Cryptococcus gattii*. Fungal Genetics and Biology 75:1–10.

Wheeler DL, Church DM, Federhen S, Lash AE, Madden TL, Pontius JU, Schuler GD, Schriml LM, Sequeira E, Tatusova TA, et al. 2003. Database resources of the National Center for Biotechnology. Nucleic Acids Research 31:28–33.

Wilson AJ, Xu J. 2012. Mitochondrial inheritance: diverse patterns and mechanisms with an emphasis on fungi. Mycology.

Wolf K, Burger G, Lang B, Kaudewitz F. 1976. Extrachromosomal inheritance in *Schizosaccharomyces pombe*. I. Evidence for an extrakaryotically inherited mutation conferring resistance to antimycin. Molecular & general genetics: MGG 144:67–73.

Wolters JF, Chiu K, Fiumera HL. 2015. Population structure of mitochondrial genomes in *Saccharomyces cerevisiae*. BMC Genomics 16:451.

Wu B, Buljic A, Hao W. 2015. Extensive horizontal transfer and homologous recombination generate highly chimeric mitochondrial genomes in yeast. Molecular Biology and Evolution 32:2559–2570.

Wu TD, Watanabe CK. 2005. GMAP: a genomic mapping and alignment program for mRNA and EST sequences. Bioinformatics 21:1859–1875.

Xiao S, Nguyen DT, Wu B, Hao W. 2017. Genetic drift and indel mutation in the evolution of yeast mitochondrial genome size. Genome Biology and Evolution 9:3088–3099.

Xu J, Ali RY, Gregory DA, Amick D, Lambert SE, Yoell HJ, Vilgalys RJ, Mitchell TG. 2000. Uniparental mitochondrial transmission in sexual crosses in *Cryptococcus neoformans*. Current microbiology 40:269–273.

Xu J, He L. 2015. Current perspectives on mitochondrial inheritance in fungi. Cell Health and Cytoskeleton 7:143–112.

Xu J, Wang P. 2015. Mitochondrial inheritance in basidiomycete fungi. Fungal Biology Reviews 29:209–219.

Yan Z, Xu J. 2003. Mitochondria are inherited from the MATa parent in crosses of the basidiomycete fungus *Cryptococcus neoformans*. Genetics 163:1315–1325.

Zhou L-W, Ji X-H, Vlasák J, Dai Y-C. 2018. Taxonomy and phylogeny of Pyrrhoderma: a redefinition, the segregation of *Fulvoderma*, gen. nov., and identifying four new species. Mycologia 26:1–18.

Zhu P, Zhai B, Lin X, Idnurm A. 2013. Congenic strains for genetic analysis of virulence traits in *Cryptococcus gattii*. Infection and immunity 81:2616–2625.

