## Supplementary Figures for "Evidence of extensive intraspecific noncoding reshuffling in a 169-kb mitochondrial genome of a basidiomycetous fungus"

**Figure S1: Genetic maps of five Hymenochaetales mitogenomes in this study.** (A) *Pyrrhoderma lamaoense,* (B) *Porodaedalea pini,* (C) *Coniferiporia sulphurascens*, (D) *Fomitiporia mediterranea* and (E) *Schizopora paradoxa*. Legends and descriptions refer to fig. 1. A *S. paradoxa* trn(gtg)-rev tRNA is encoded on negative strand but plotted at outer circle for aesthetic reasons.

(A)

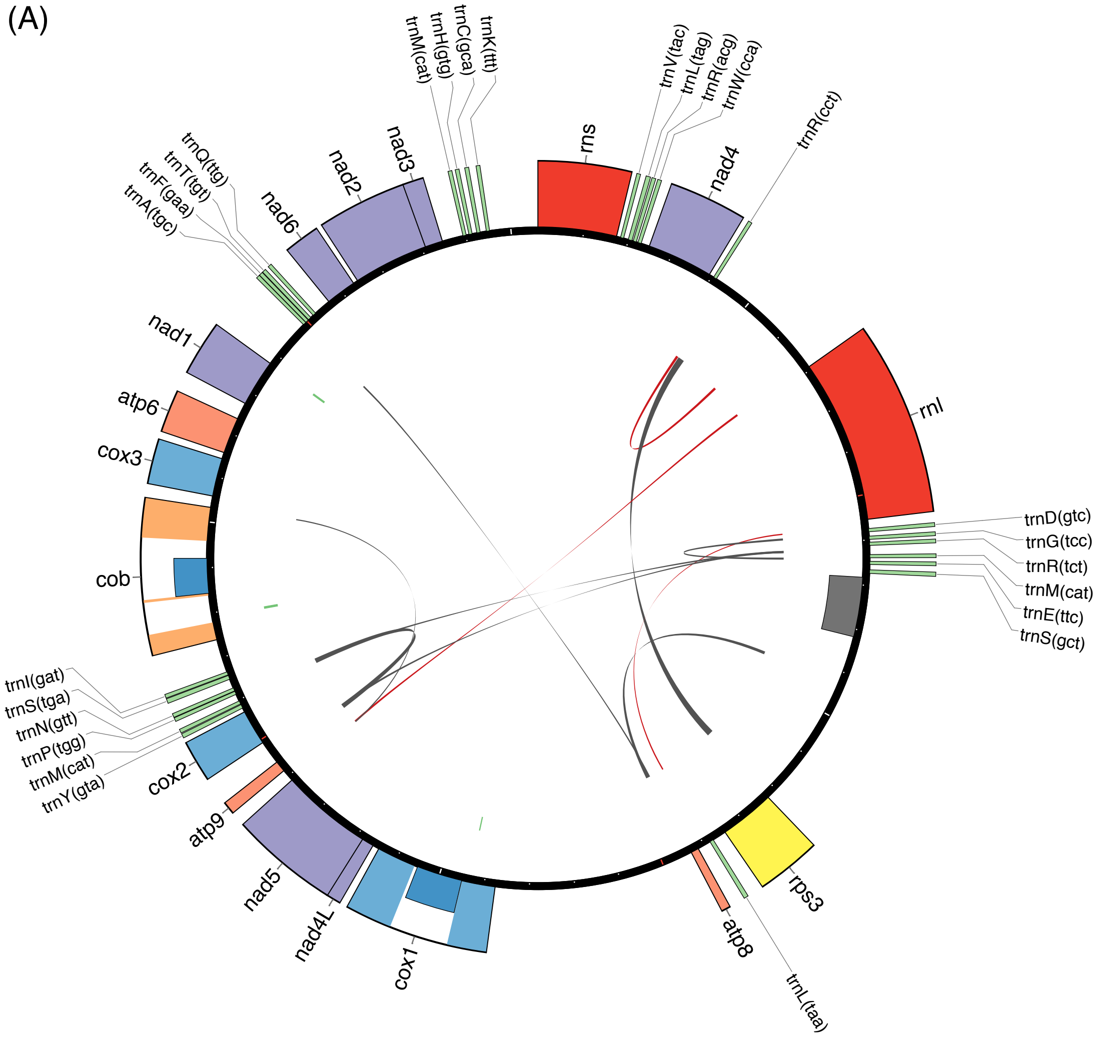

(B)

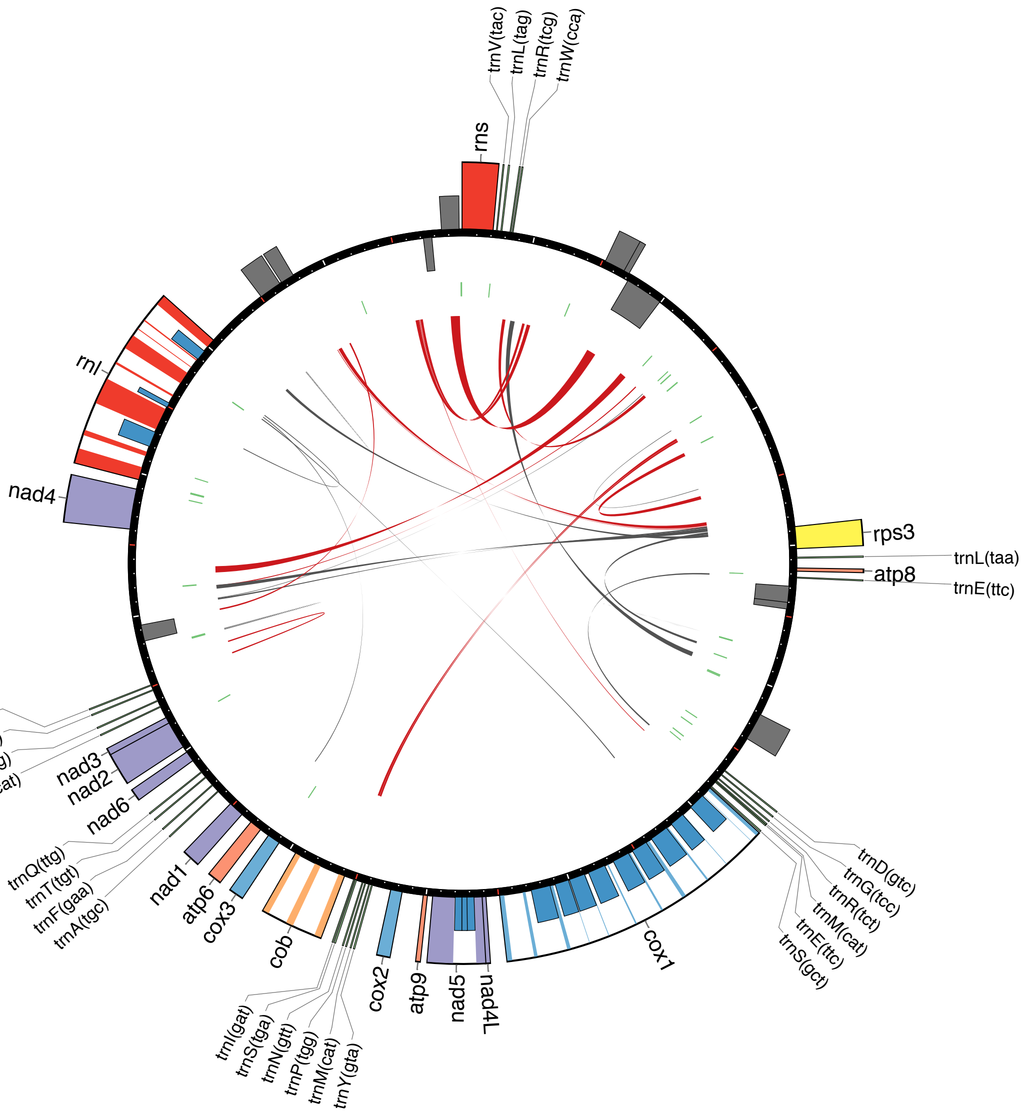
trnV(tac)
trnL(tag)
trnR(tcg)
trnW(cca)
rns
rnl
nad4
rps3
trnL(taa)
atp8
trnE(ttc)
trnK(ttt)
trnC(gca)
trnH(gtg)
trnM(cat)
nad3
nad2
nad6
trnQ(ttg)
trnT(tgt)
trnF(gaa)
trnA(tgc)
nad1
atp6
cox3
cob
trnI(gat)
trnS(tga)
trnN(gtt)
trnP(tgg)
trnM(cat)
trnY(gta)
cox2
atp9
nad5
nad4L
cox1
trnD(gtc)
trnG(tcc)
trnR(tct)
trnM(cat)
trnE(ttc)
trnS(gct)

(C)

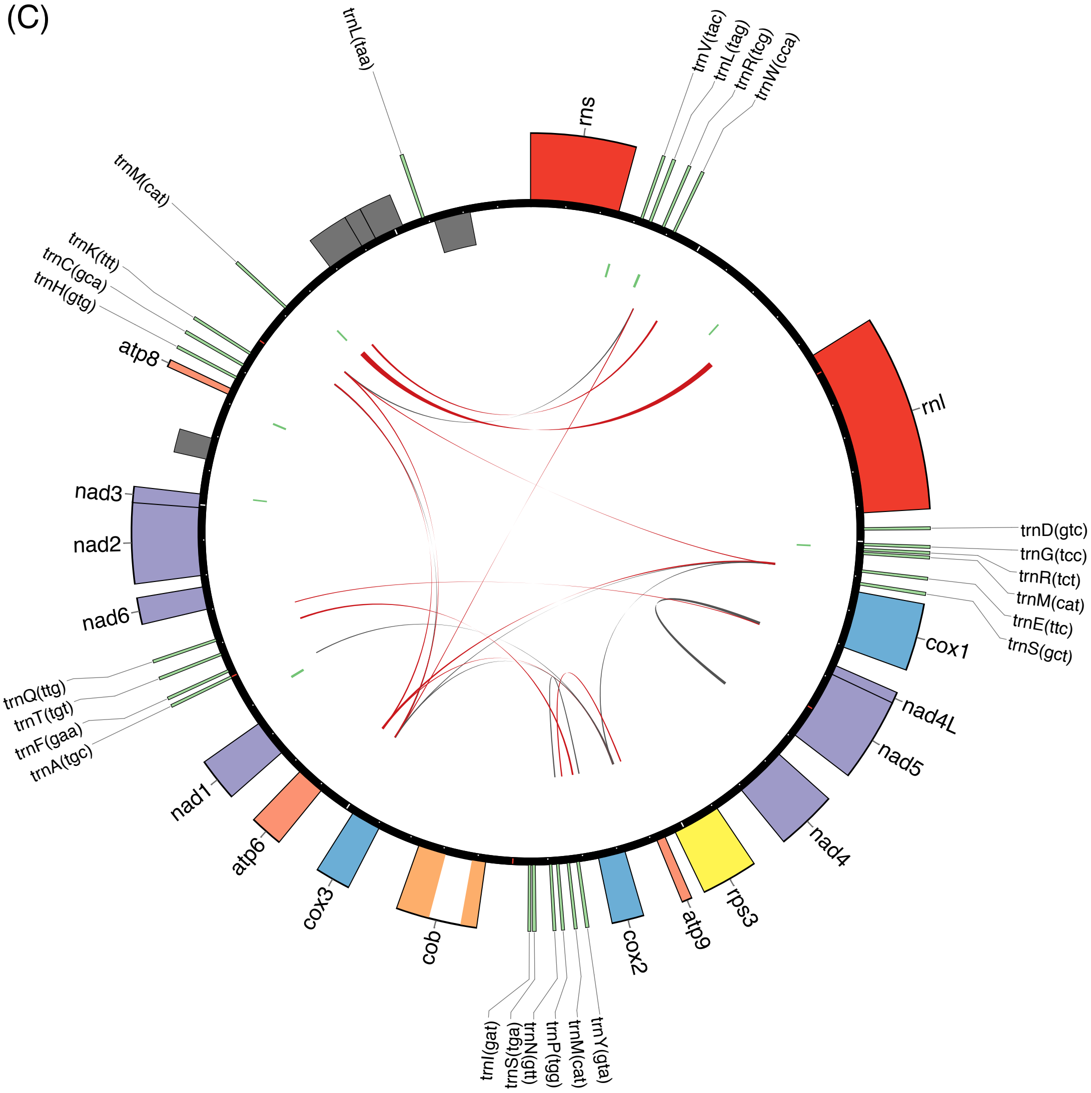

trnL(taa)
rns
trnV(tac)
trnL(tag)
trnR(tcg)
trnW(cca)
trnM(cat)
trnK(ttt)
trnC(gca)
trnH(gtg)
atp8
rnl
nad3
nad2
trnD(gtc)
trnG(tcc)
trnR(tct)
trnM(cat)
trnE(ttc)
trnS(gct)
nad6
cox1
trnQ(ttg)
trnT(tgt)
trnF(gaa)
trnA(tgc)
nad4L
nad5
nad1
nad4
atp6
cox3
rps3
atp9
cob
cox2
trnI(gat)
trnS(tga)
trnN(gtt)
trnP(tgg)
trnM(cat)
trnY(gta)

(D)

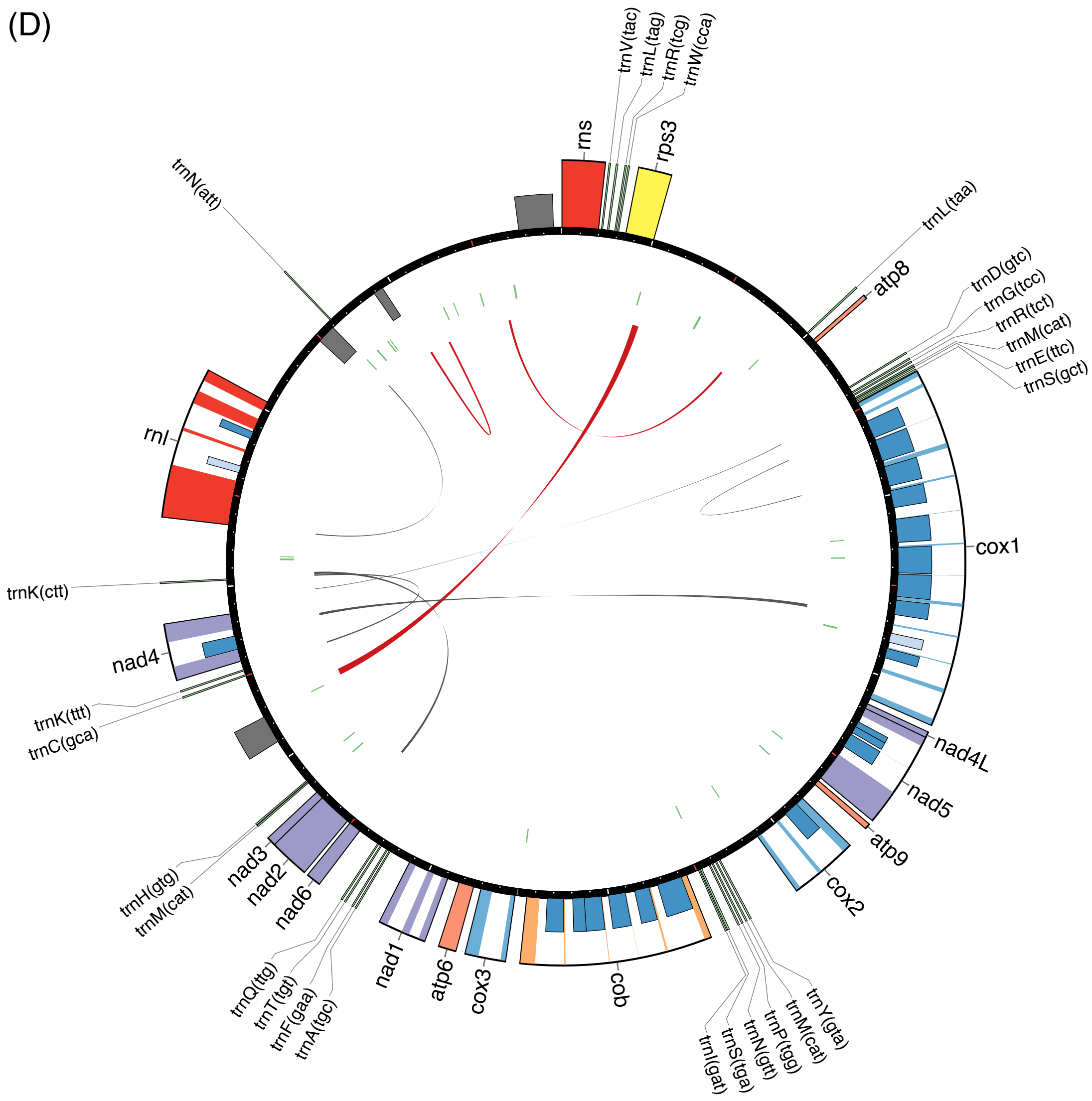

rns
trnV(tac)
trnL(tag)
trnR(tcg)
trnW(cca)
rps3
trnL(taa)
atp8
trnD(gtc)
trnG(tcc)
trnR(tct)
trnM(cat)
trnE(ttc)
trnS(gct)
cox1
nad4L
nad5
atp9
cox2
trnY(gta)
trnM(cat)
trnP(tgg)
trnN(gtt)
trnS(tga)
trnI(gat)
cob
cox3
atp6
nad1
trnA(tgc)
trnF(gaa)
trnT(tgt)
trnQ(ttg)
nad6
nad2
nad3
trnM(cat)
trnH(gtg)
trnC(gca)
trnK(ttt)
nad4
trnK(ctt)
rnl
trnN(att)

(E)

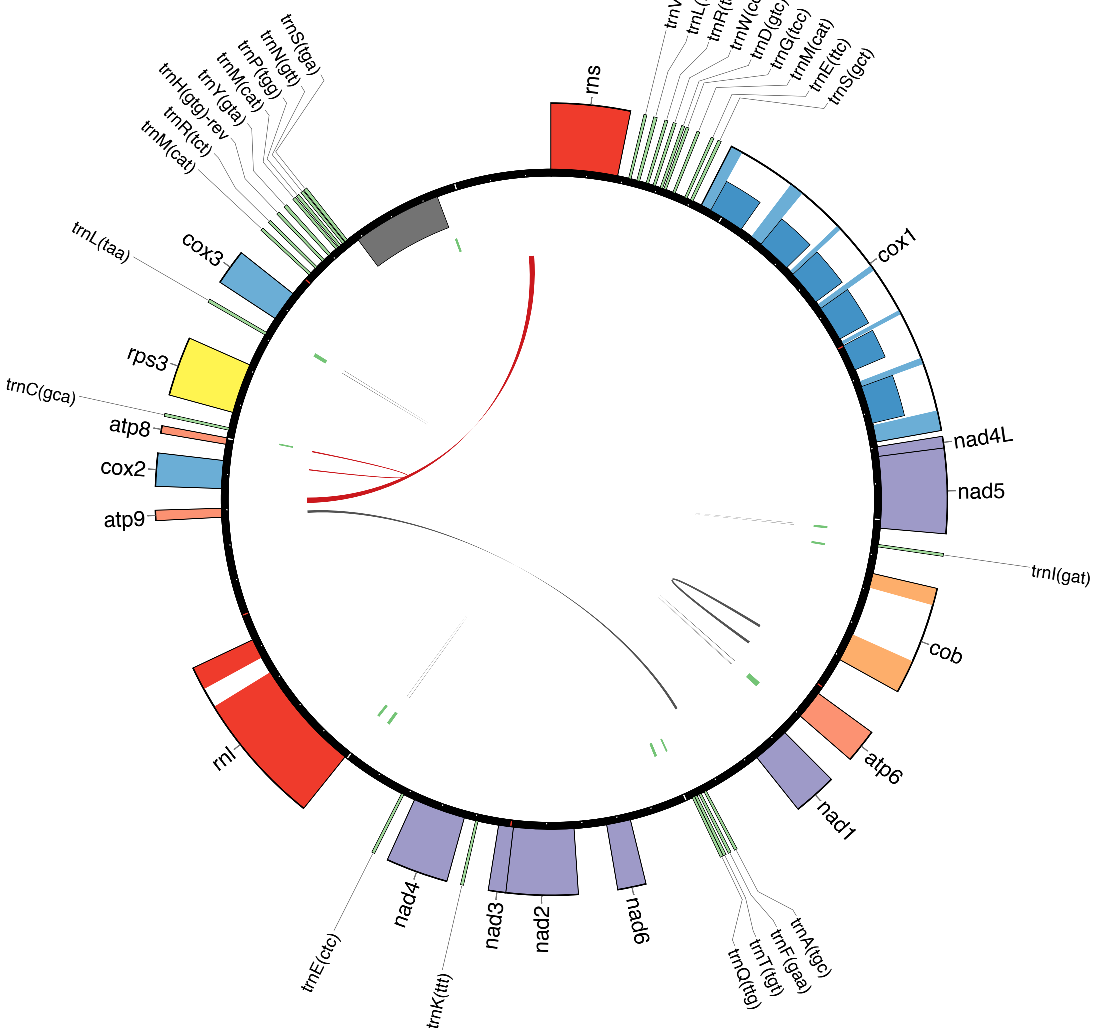

rns
trnV(tac)
trnL(tag)
trnR(tcg)
trnW(cca)
trnD(gtc)
trnG(tcc)
trnM(cat)
trnE(ttc)
trnS(gct)
cox1
nad4L
nad5
trnI(gat)
cob
atp6
nad1
trnA(tgc)
trnF(gaa)
trnT(tgt)
trnQ(ttg)
nad6
nad2
nad3
trnK(ttt)
nad4
trnE(ctc)
rnl
atp9
cox2
atp8
trnC(gca)
rps3
trnL(taa)
cox3
trnM(cat)
trnR(tct)
trnH(gtg)-rev
trnY(gta)
trnM(cat)
trnP(tgg)
trnN(gtt)
trnS(tga)

**Figure S2: Scaled plots of mitogenome synteny of (A) six Hymenochaetales species and (B) comparation between *Pyrrhoderma noxium* and *Pyrrhoderma lamaoense*.** Red arrows are mitochondrial core genes and green tiles are tRNA genes. The red links between two mitogenomes are aligned regions with identity > 80 %. The plot was generated with genoplotR package in R.

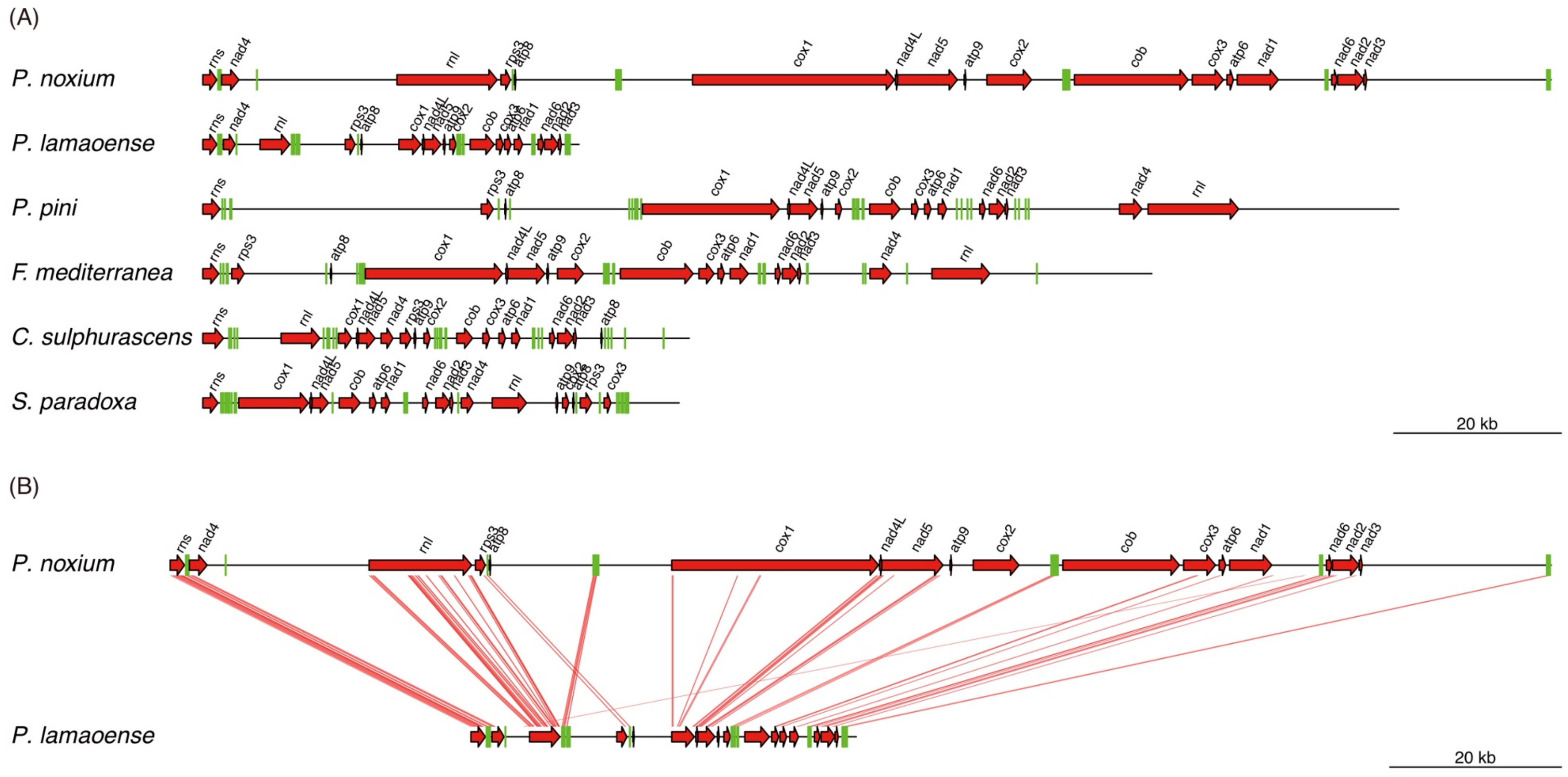

**Figure S3: Intron phylogeny of 3 intron sites shared by more than 4 species.** The bootstrap values were inferred with 100 replicates.

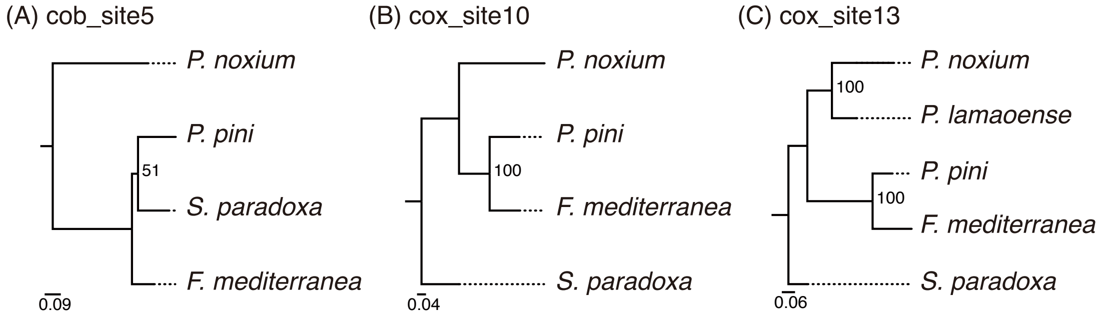

**Figure S4: Alleles of amino acid comparing to KPN91.** The x labels show gene names and positions in align protein sequences. The texts in plot are amino acid abbreviations.

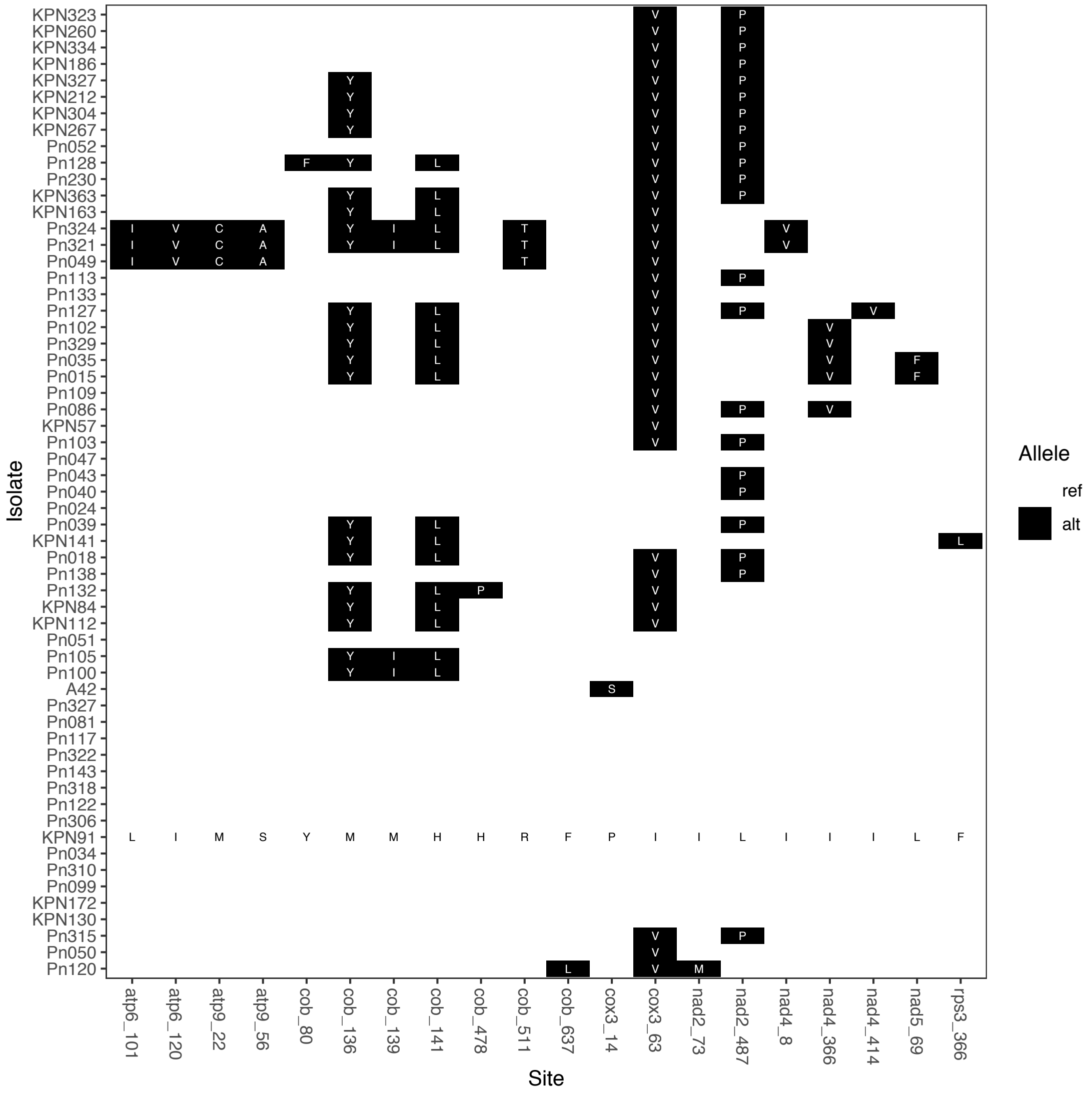

**Figure S5: Length of all introns in 59 *Pyrrhoderma noxium* isolates.** Length is normalized to KPN91. Blank tiles indicate lack of intron.

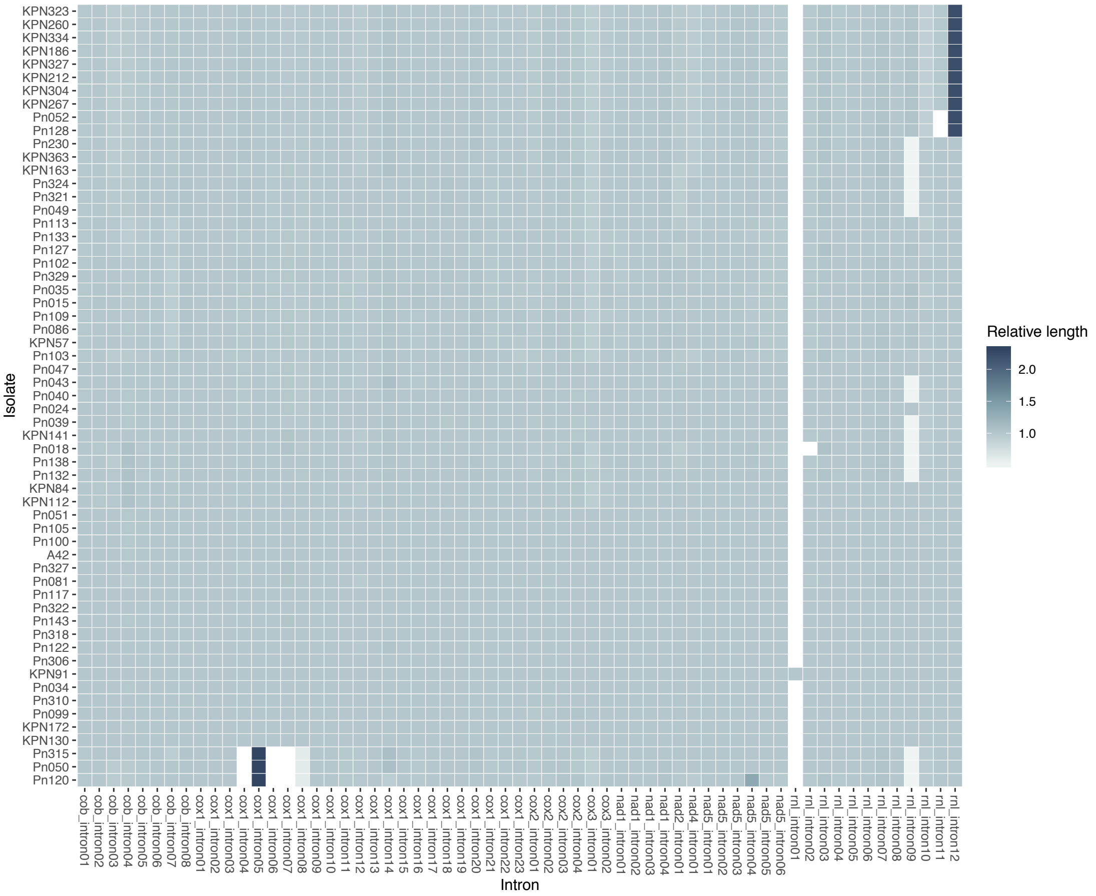

**Figure S6: Phylogenetic tree of concatenated intron sequences.** Red dots represent bootstrap value > 80. Tiles are the lineages defined by intronic phylogeny and intron pattern.

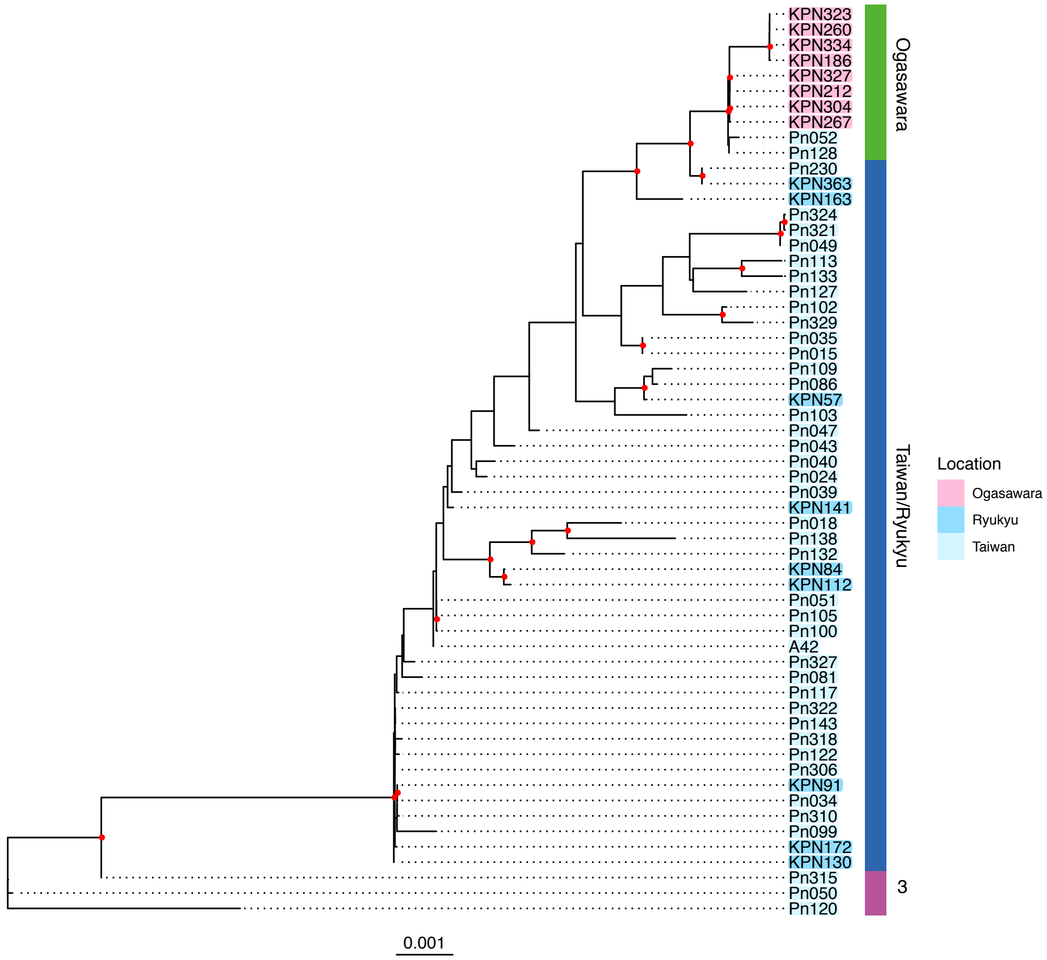

**Figure S7: Example mitogenome synteny plots showing the rearrangement events.** Legends refer to figure S2. (A) Intra- and inter-rearrangements in intergenes between KPN91 (Taiwan/Ryukyu lineage) and KPN323 (Ogasawara lineage). (B) No rearrangement between KPN91 (Taiwan/Ryukyu lineage) and KPN130 (Taiwan/Ryukyu lineage). The only gap is an exclusive *rnl* intron in KPN91. (C-D) Rearrangement and INDEL result in alternation of a tRNA (black arrow).

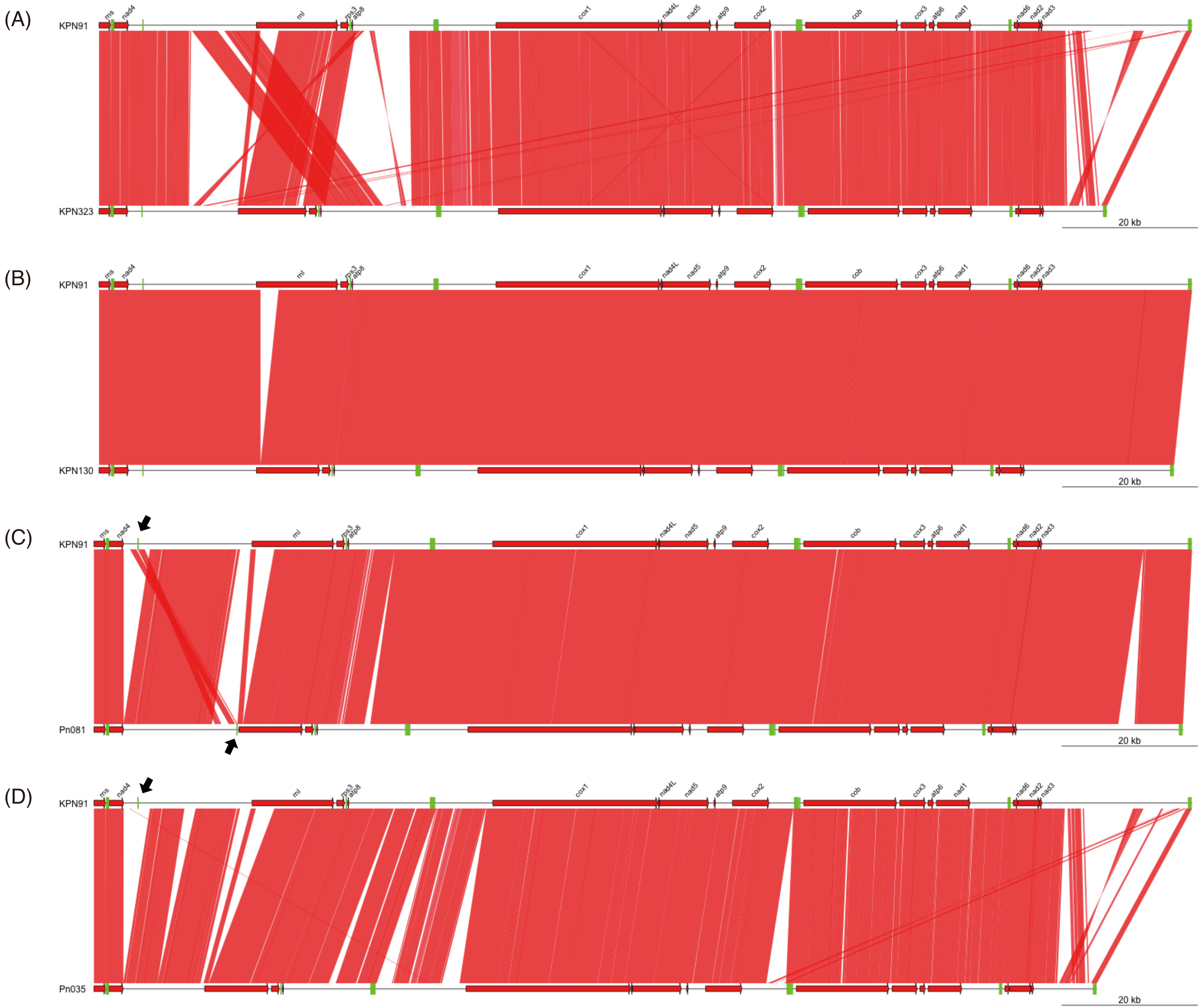
(A)
KPN91
KPN323
20 kb
(B)
KPN91
KPN130
20 kb
(C)
KPN91
Pn081
20 kb
(D)
KPN91
Pn035
20 kb
rns
nad4
rnl
rps3
atp8
cox1
nad4L
nad5
atp9
cox2
cob
cox3
atp6
nad1
nad6
nad2
nad3

**Figure S8: Classification of intergene rearrangement events against KPN91 (Taiwan/Ryukyu lineage) and KPN323 (Ogasawara lineage).** A rearrangement event is defined according to examples in figure S7. Prefixes of x labels represent intra- and inter-rearrangement, respectively; suffixed represent intergenic regions defined in fig. 1. No: no rearrangement against reference.

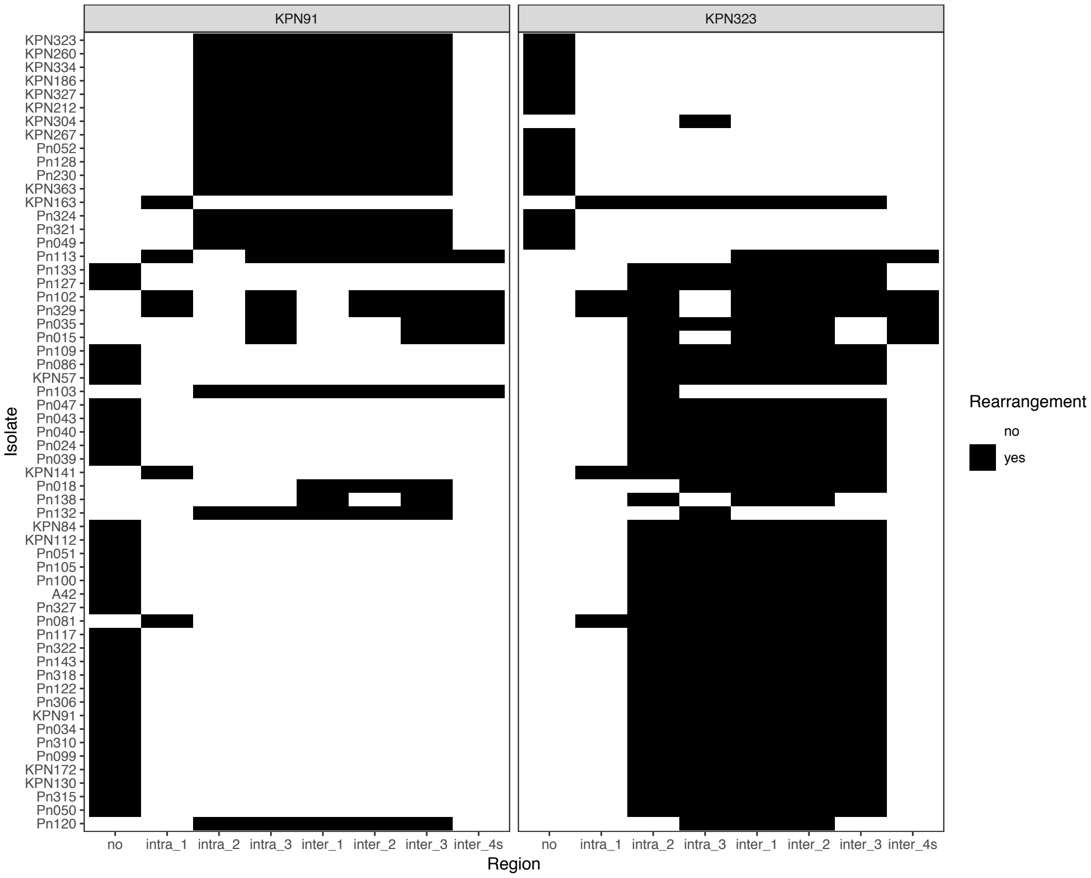
